# Systematic mapping of bacteriophage gene essentiality with HIDEN-SEQ

**DOI:** 10.1101/2025.11.20.689424

**Authors:** Dorentina Humolli, Jessica Ransome, Damien Piel, Jan-Willem Veening, Alexander Harms

## Abstract

The constant arms race of bacteriophages and their bacterial hosts has inspired major breakthroughs in biotechnology and shaped phages as fierce predators with great clinical potential to fight multidrug-resistant bacterial pathogens. However, the vast amount of genomic “dark matter” composed of genes of unknown function in phage genomes remains a major obstacle for the molecular understanding of phage-host interactions. Here we present HIDEN-SEQ, a transposon-insertion sequencing method for phages that systematically links viral genes to selectable phenotypes. Using model phage T4, we show that HIDEN-SEQ readily reproduces the gene essentiality map established over decades of research. Subsequently, we show that our method is easily portable to different phages far beyond classical laboratory models. Across a panel of bacterial hosts and growth conditions, HIDEN-SEQ reveals many conditionally essential phage genes, including previously unknown viral anti-defense factors that we could match to specific antiviral defenses of the respective hosts. Compared to analogous techniques, HIDEN-SEQ provides unprecedented depth and near base-pair resolution as well as great ease of use and portability. We therefore anticipate that HIDEN-SEQ will accelerate discoveries in phage biology by uncovering functions of viral dark matter with direct relevance for microbial ecology, biotechnology, and improvements of phage therapy.

## Main

Bacteriophages, the viruses infecting bacteria, are the most abundant and genetically diverse biological entities on Earth^1,2^. Over billions of years, these viral predators and their hosts have co-evolved in a dynamic arms race that is constantly shaping microbial communities, balancing ecosystems, and driving biogeochemical cycles around the globe^3–5^. Research on phage-host interactions using a few model phages has been crucial for our understanding of the basic molecular biology of life, e.g., by unraveling the nature of the genetic code, and has provided the tools for critical technological breakthroughs such as molecular cloning^6^. Furthermore, the current crisis of antimicrobial therapy due to increasing rates of multidrug resistance is driving a renaissance of clinical applications of phages to combat bacterial infections (“phage therapy”)^7^. However, a major limiting factor for phage-inspired biotechnology and rational applications of phage therapy remains the vast genetic “dark matter” of enigmatic genes with no known biological function that dominate phage genomes^2,8^. We have therefore so far just scratched the surface of the true potential of these viruses to understand ecology, drive innovation in biotechnology, and fight antibiotic-resistant infections in clinics.

Most tailed phages have genomes of 50-200 kb in size of which one part is composed of operons of core genes encoding functions involved in phage replication, gene expression, and virion assembly^2,9^. The other genes, largely coding for hypothetical proteins, are often dispensable under standard laboratory conditions and their proportion increases with genome size^9–11^. Based on few previous studies, these accessory genes provide functions that enhance the ability of the phage to survive and adapt to specific hosts or environmental conditions^11,12^. As an example, phage T7 encodes a serine-threonine kinase (Gp0.7) which targets various bacterial proteins to enable growth on starved hosts^13^ and to interfere with bacterial immunity^14,15^. Other examples are T4 genes encoding inhibitors of specific antiviral defense systems like *rIIA*/*rIIB* for RexAB or *dmd* for RnlAB and various anti-CRISPR proteins encoded in different phages^16,17^. Accessory genes of bacteriophages thus seem to largely form their arsenal of tools and tricks to prevail in the virus-host arms race which makes them a crucial reservoir for innovation in biotechnology and for improvements of phage therapy. To systematically uncover the biological functions of these genes, scalable genome-wide approaches linking viral genes to specific phenotypes are required. Existing methods have offered valuable insights but remain laborious and not broadly applicable^18–23^. In bacteria, this challenge has been overcome by transposon-insertion sequencing (TnSeq), a technique that quantifies the fitness of all viable transposon insertions in a genome by comparative deep sequencing before and after phenotypic selection^24^. While TnSeq has revolutionized bacterial microbiology^25^, no analogous approach exists for phages. Although transposition into phage genomes had occasionally been reported^26–28^, its scale and application have been limited by the lack of efficient genetic selection markers.

Here we present HIDEN-SEQ, a functional and easy-to-use TnSeq system for phages that combines highly efficient transposition into viral genomes with CRISPR-based selection for phages that have acquired a transposon. Using HIDEN-SEQ, we have built essentiality maps at unprecedented near base-pair resolution for three different phages and demonstrate its utility for the identification of conditionally essential genes, including diverse new *bona fide* anti-defense factors and genes that are crucial under specific growth conditions. We further applied HIDEN-SEQ to probe the conditional essentiality of phage genes across a panel of bacterial strains from clinical infections which revealed highly specific mechanisms of how different phages interact with the bacterial cell envelope and multiple layers of bacterial immunity.

### HIDEN-SEQ enables transposon-insertion sequencing in phages

We began our study by setting the basics of TnSeq for bacteriophages using the well-studied model phage T4 infecting laboratory organism *Escherichia coli* K-12. For transposition, we built a new system based on *mariner* transposon Himar1 that is commonly used for TnSeq^29^ and inserts randomly at TA dinucleotide sites (see details in *Methods*). The Himar1 transposon was engineered to encode the anti-CRISPR gene *acrVIA1*^30^. For selection, we expressed LseCas13a nuclease in the host with an efficient crRNA targeting an essential T4 gene (Extended Data Fig. 1a). Only phage clones that acquire the transposon, and thus express AcrVIA1, can evade Cas13a targeting and replicate^31^. This variant of CRISPR-Cas is well suited for selection of transposition across different phages because it targets RNA^32^, not DNA (inherently bypassing viral genome protection), and it restricts most phages that have been tested so far^31,33^. We call this setup HIDEN-SEQ (<u>hid</u>den Acr-<u>en</u>abled transposon-insertion <u>seq</u>uencing) because host-expressed CRISPR-Cas13a and the transposon-encoded Acr protein of the phage play “hide and seek” in infected cells (Fig. 1a).

**Fig. 1.**
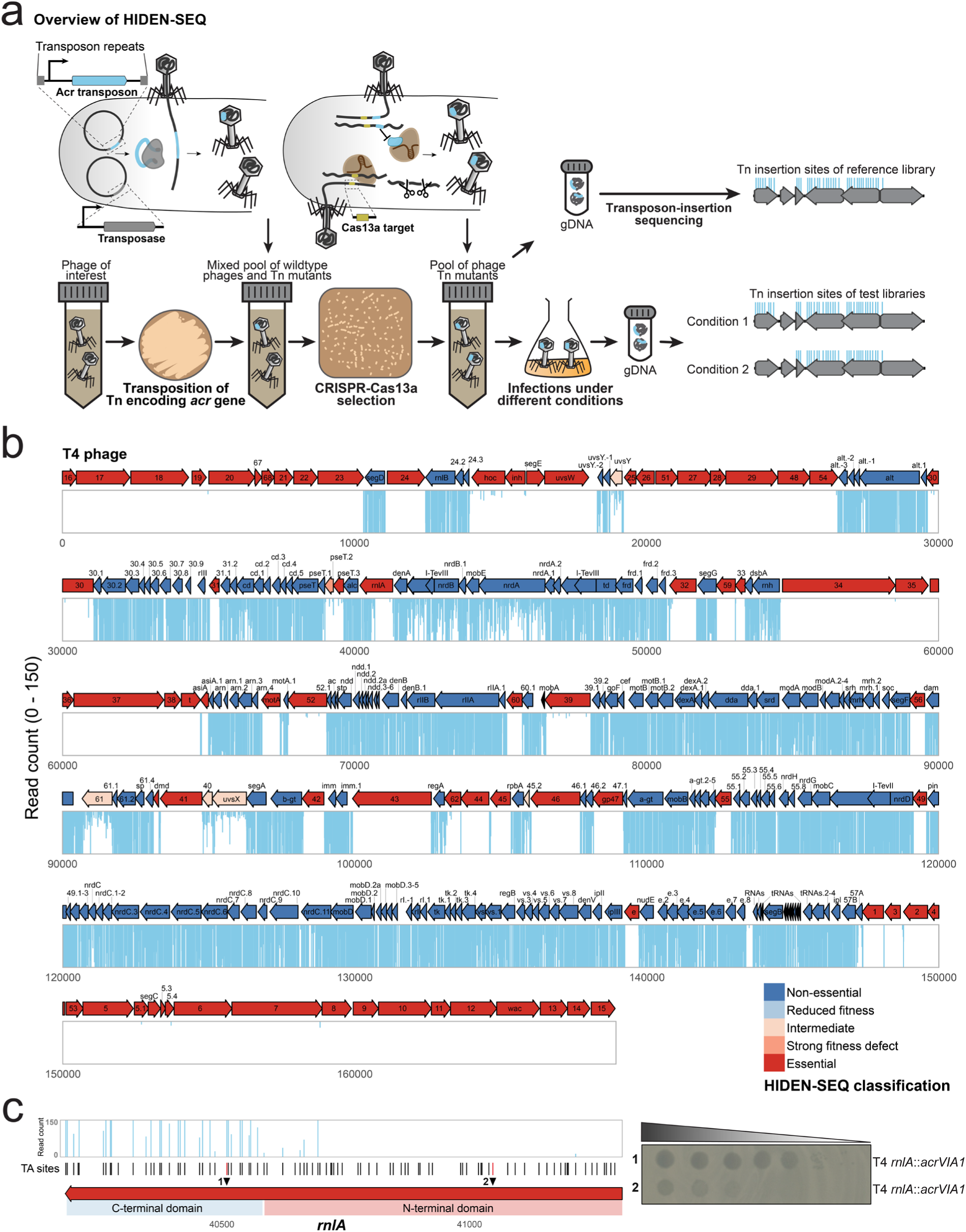
Combining transposition and CRISPR-Cas13a based selection as HIDEN-SEQ for phage T4. **a**, Transposition introduces an anti-CRISPR gene (*acr*) randomly into TA dinucleotide sites within the phage genome. *acr* expression inhibits CRISPR-Cas13a immunity and enables the selection of a library of phage transposon mutants. Deep sequencing of insertion sites is performed to quantify the abundance of transposon mutants before (reference) and after exposing the library to different conditions. **b**, Phage T4 HIDEN-SEQ gene essentiality map with position 1 set to the start of the small terminase subunit gene. Arrows present the coding sequences with the orientation indicating the direction of transcription. Only the full-length gene products are shown. Arrow colours correspond to HIDEN-SEQ classification categories derived from transposon insertion profiles: non-essential (dark blue), reduced fitness (light blue), intermediate (light orange), strong fitness defect (orange), and essential (red). Vertical bars represent the number of transposon insertions at TA dinucleotide sites with heights indicating read counts (up to a maximum of 150). Genes containing ≤5 TA sites are shown in black and were not classified. **c**, (Left) Distribution of transposon insertions across T4 *rnlA* gene with C-terminal and N-terminal domains indicated. Two TA sites disrupted in mutants used for validation are marked as site 1 and site 2. (Right) Ten-fold serial dilution plaque assays of the T4 engineered mutants carrying insertions at sites 1 and 2 on the *E. coli* K-12 ΔRM strain, confirming the domain essentiality pattern observed in the HIDEN-SEQ data.

To assess whether transposition occurs efficiently in the phage genome and if these mutants can be selectively enriched, we plated T4 which had been targeted with the anti-CRISPR transposon or a non-protective control on the selection host (Extended Data Fig. 1b). As expected, CRISPR-Cas13a selection completely abolished T4 growth but many viral clones were obtained after transposition with the anti-CRISPR transposon. Whole-genome sequencing of thirteen individual clones confirmed that all carried unique transposon insertions (Extended Data Fig. 1c).

Subsequently, we applied HIDEN-SEQ to build a genome-wide gene essentiality map of model phage T4 to test the potential of our technique. This phage has a 168 kb genome encoding nearly 300 genes of which around 130 have remained uncharacterized despite decades of research^10^. The T4 input HIDEN-SEQ library, collected before CRISPR-Cas13a selection, showed abundant insertions distributed across the genome (Extended Data Fig. 2a, b), confirming efficient transposition. Reads in essential genes are expected at this step because the library contains mutants from the initial viral burst after transposition with minimal subsequent replication. After outgrowth, our results show that 190 genes in the T4 genome are not essential under standard laboratory conditions which is well in agreement with the literature^10^ (Fig. 1b and Extended Data Fig. 2c). Importantly, the unprecedented depth and resolution enabled by HIDEN-SEQ allowed us to extend the essentiality classification beyond binary calls to include weakly, moderately, and strongly reduced fitness of gene disruptions (see *Methods*). As an example, five genes in the T4 genome (*40*, *45.2*, *61*, *uvsX*, *uvsY*) showed a low but still detectable tolerance to transposon insertions, indicating that their disruption imposed a fitness defect but did not completely abolish phage viability. Furthermore, HIDEN-SEQ enables sub-gene resolution of fitness analyses as illustrated by T4 RNA ligase A (*rnlA)*^34^, which tolerates insertions in its C-terminal but not in the N-terminal catalytic domain (Fig. 1c). Taken together, these results establish HIDEN-SEQ as an effective tool for mapping phage gene essentiality at unprecedented depth and resolution.

### HIDEN-SEQ identifies known and novel conditionally essential genes in phage T4

To test whether HIDEN-SEQ can identify conditional essential genes, we used our highly saturated T4 HIDEN-SEQ library to infect hosts expressing various bacterial defense systems. This allowed us to directly evaluate whether insertions in known anti-defense genes, which are non-essential under standard conditions, would be selectively depleted when challenged by the targeted defense system. We first examined the *rIIA* and *rIIB* genes of T4 which counteract the RexAB defense system^16^ that protects lambda lysogens from other phages by abortive infection due to membrane depolarization^35^. As expected, transposon insertions in *rIIA* and *rIIB* genes were significantly depleted when the T4 library was grown in presence of RexAB while both genes were non-essential without it (Fig. 2a and Extended Data Fig. 3a). Subsequently, we focused on *dmd* which encodes an inhibitor of the antiviral RnlAB toxin-antitoxin system^36^. Our initial experiments had identified the *dmd* gene of T4 phage as essential because the *E. coli* K-12 laboratory strain naturally expresses RnlAB^36^ (Fig. 1b). We thus generated another T4 transposon library in a *rnlA* knockout strain as host and used it to infect this mutant and the parental wildtype host. As expected, transposon insertions in *dmd* were strongly depleted in the host expressing RnlAB but not in the knockout control (Fig. 2b and Extended Data Fig. 3a). Importantly, when comparing the two independently generated T4 libraries, no other gene showed significant differences, highlighting both the specificity of bacterial immunity and the reproducibility of HIDEN-SEQ (Extended Data Fig. 3b).

**Fig. 2.**
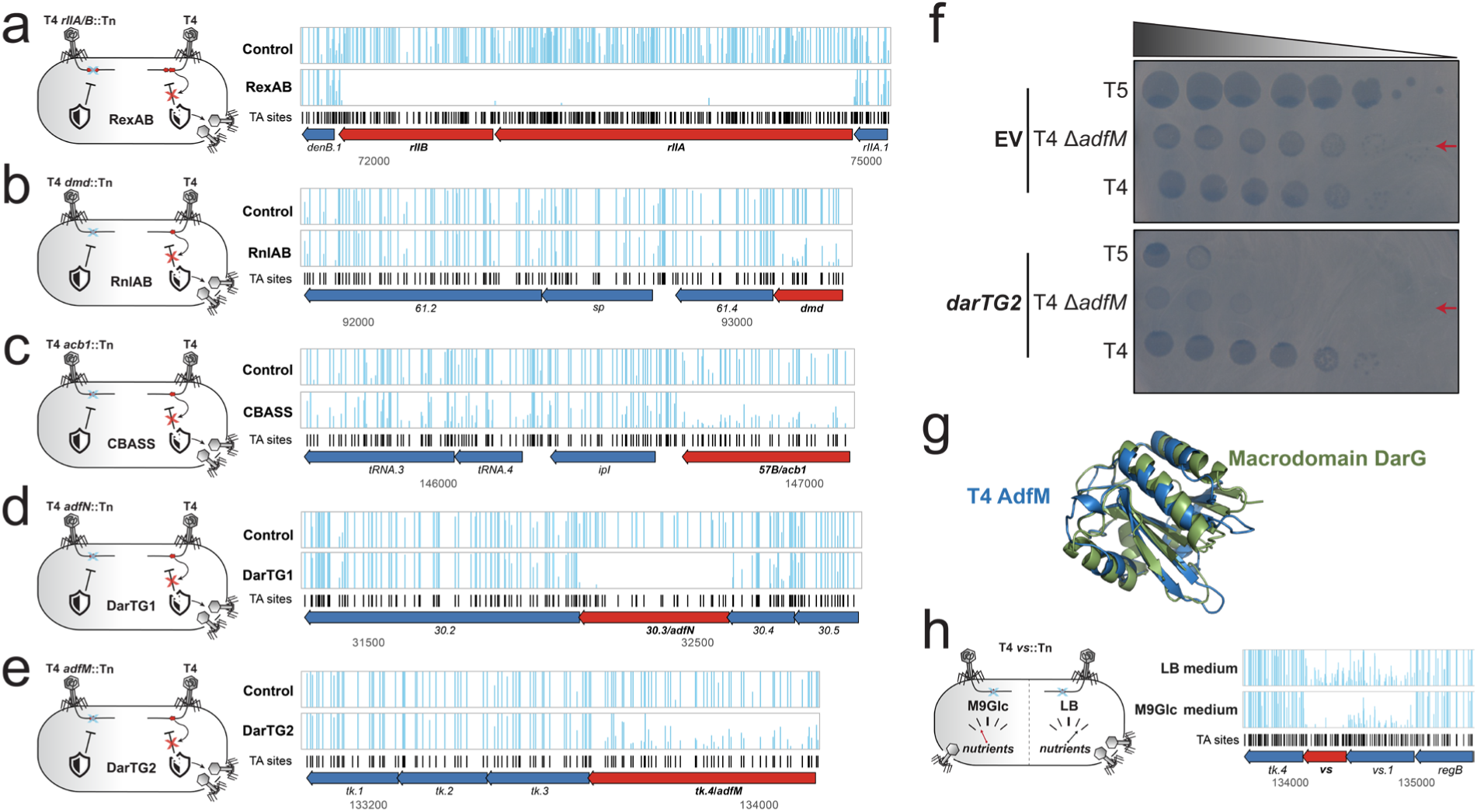
Identification of conditionally essential phage genes by HIDEN-SEQ. The T4 phage reference HIDEN-SEQ library was used to infect different variants of the *E. coli* K-12 BW25113 host strain to compare viral gene essentiality in the presence of different anti-phage defense systems. For each panel (**a-e**), transposon insertion profiles obtained after infection in the presence of the defense system condition (bottom track) are compared with the respective control (top track). Vertical bars represent the number of transposon insertions at TA sites (shown with black tick marks) with heights indicating read counts (up to a maximum of 150). Arrows indicate coding sequences with orientation showing direction of transcription. Genes highlighted in red were identified as conditionally essential in the presence of the indicated defense system and are shown with their surrounding genomic regions. **a**, empty vector control compared to *E. coli* K-12 expressing *rexAB*. **b**, *E. coli* K-12 Δ*rnlA* compared to *E. coli* K-12 expressing *rnlAB.* **c**, *E. coli* K-12 carrying a plasmid encoding a catalytically inactive (control) or active CBASS operon. **d**, empty vector control compared with *E. coli* K-12 expressing *darTG1*. **e**, empty vector control compared with *E. coli* K-12 expressing *darTG2*. **f**, Ten-fold serial dilution plaque assays on *E. coli* K-12 carrying an empty vector (EV) control or *darTG2* expression plasmid challenged with phage T5 (sensitive to DarTG2), phage T4 engineered to remove AdfM (Δ*adfM*), or wildtype phage T4. Data are representative of 3 independent biological replicates. **g**, Structural alignment of AlphaFold2 predicted structures of T4 AdfM with the macrodomain of *Mycobacterium tuberculosis* DarG homolog (probability = 99.71%, E-value = 2.6 × 10⁻¹⁴). **h**, Transposon insertion profiles obtained after infection of *E. coli* K-12 BW25113 with T4 reference HIDEN-SEQ library in M9Glc medium (bottom track) and LB medium (top track), highlighting in red the strongest depleted *vs* gene in the M9Glc condition. Vertical bars represent the number of transposon insertions at TA sites (shown with black tick marks) with heights indicating read counts (up to a maximum of 150).

In addition, T4 phage encodes an anti-CBASS protein (57B/Acb1) which degrades the cyclic nucleotides generated by this defense system as a second messenger in response to phage infection^37^. As expected, *57B/acb1* appeared as non-essential in our reference transposon library grown on *E. coli* K-12 which lacks CBASS. However, when we challenged this library with a host expressing CBASS from a plasmid^37^, we observed a clear reduction in read counts across this gene (Fig. 2c). Finally, we used HIDEN-SEQ to probe the biology of an anti-DarTG gene of phage T4, *30.3/adfN*, that encodes an enzyme reverting the ADP-ribosylation of DNA which is added by the DarT toxin in response to phage infection to block viral replication^38^. As expected from previous work^38^, *30.3/adfN* was found to be not essential for growth on *E. coli* K-12, but insertions in this gene were completely lost in presence of DarTG1 defense system (Fig. 2d and Extended Data Fig. 3a).

While *adfN* protects T4 against DarTG1, it does not act against DarTG2, probably because NADAR proteins like AdfN act on modified guanosine bases, whereas DarTG2 modifies thymidines^38^. Since T4 lacking *adfN* remains resistant to DarTG2^38^, we suspected that T4 may additionally encode an anti-DarTG2 factor. To test this, we used the T4 HIDEN-SEQ library to infect a host expressing DarTG2 and found a single significant hit, *tk.4*, which showed reduced read counts specifically in presence of this defense system (Fig. 2e). Consistently, a clean knockout of *tk.4* sensitized phage T4 to DarTG2 (Fig. 2f). Interestingly, Tk.4 shows structural similarity to the DarG antitoxin of *Mycobacterium tuberculosis* DarTG2^39^, and both proteins are homologs featuring macrodomains which typically act as an ADP-ribosylglycohydrolase^40^ (Fig. 2g). Based on these findings, we conclude that *tk.4* encodes a previously unknown anti-DarTG2 factor, which we name anti-DarT factor Macrodomain (*adfM*). Analogous to AdfN^38^, it seems likely that AdfM neutralizes DarT2 toxicity by enzymatic removal of ADP-ribose modifications from the DNA.

To test whether conditional essential genes can also be identified beyond the context of bacterial immunity, we compared the mutant fitness of the T4 HIDEN-SEQ library in rich LB medium and defined M9 minimal medium (see *Methods*). Under nutrient limitation, where host and phage need to synthesize all biomolecules *de novo* from a handful of compounds, the strongest depletion was observed for the T4 *vs* gene encoding a modifier of valyl-tRNA synthetase (Fig. 2h and Extended Data Fig. 3c). While the biological role of this modification is not known^41^, deletion of *vs* impaired T4 plaque formation specifically in M9Glc medium (Extended Data Fig. 3d), confirming a fitness defect dependent on nutrient availability.

Taken together, these results demonstrate that HIDEN-SEQ can identify conditionally essential genes, including known and novel factors required by phages to overcome bacterial defense systems and those enabling phage replication at different physiological states of the host.

### HIDEN-SEQ is applicable across different *Escherichia coli* phages

To explore the potential of HIDEN-SEQ beyond model phage T4, we next applied it to other phages infecting *E. coli*. First, we constructed a transposon mutant library of phage Bas37 (Extended Data Fig. 4a-c), a close relative of T4^42^, which allowed us to evaluate the robustness of HIDEN-SEQ results by comparing gene essentiality between related phages. Overall, Bas37 and T4 genomes showed highly similar gene essentiality patterns with few minor differences (Fig. 3a), mostly due to technicalities of the read count distribution and classification (Extended Data Fig. 4d). Beyond these, the RNA ligase A of Bas37 (*bas37_0061*) showed essentiality across both protein domains, whereas in T4 only the N-terminal domain appeared essential (Fig. 3a). Furthermore, we found that *bas37_0161*, the ortholog of *regA*, a translational repressor of early phage genes, was essential in Bas37 but non-essential in T4 under standard laboratory conditions (Fig. 3a). These results reveal specific differences in the interaction of T4 and Bas37 with *E. coli* K-12 which could, e.g., be informative about different interactions with bacterial immunity of this host.

**Fig. 3.**
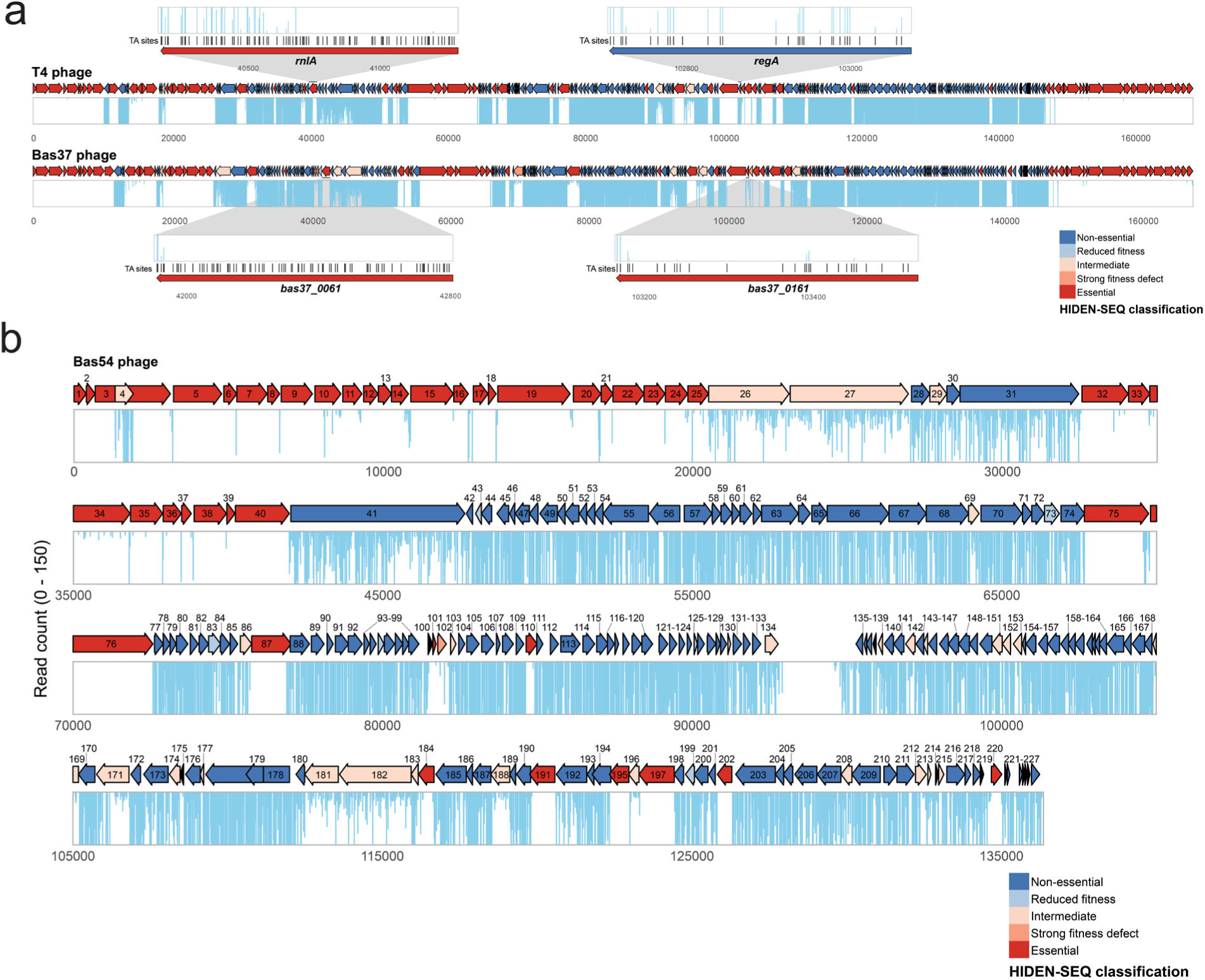
Genome-wide essentiality maps of different *E. coli* phages. **a**, HIDEN-SEQ essentiality maps of T4 and Bas37, two closely related phages, showing broadly similar transposon insertion patterns across both genomes. Zoomed-in views highlight two genes in phage Bas37 that show different insertion patterns compared to their orthologs in phage T4. **b**, Phage Bas54 HIDEN-SEQ gene essentiality map with position 1 set to the start of the i-spanin gene at the beginning of the terminase operon. Colour coding as in Fig. 1b.

After comparing these two T-even phages, we applied HIDEN-SEQ to Bas54, a phage of the poorly studied *Vequintavirinae* subfamily (“V5-like phages”) with a genome size of 136 kb encoding 227 predicted genes^42^. Using this phage, we also tested the flexibility of our method by using a different variant of CRISPR-Cas13a, the LbuCas13a system^33^, during selection of the transposon mutants (Extended Data Fig. 5a) expressing AIcrVIA3, a novel AI-designed efficient inhibitor of the LbuCas13a nuclease^43^. Due to an overestimation of mutants during pooling, our initial Bas54 library was not sufficiently saturated, limiting the resolution of essentiality calls (Extended Data Fig. 5b, c). We therefore repeated the library construction by collecting a larger pool of mutants, which resulted in a highly saturated library with much clearer biological results (Fig. 3b and Extended Data Fig. 5d). Our data show that 146 genes in the Bas54 genome are not essential under standard laboratory conditions, including 6 genes where disruptions caused markedly reduced fitness (Fig. 3b). Like other *Vequintavirinae* and related phages, Bas54 encodes an extensive set of predicted tail fibers, which has been suggested to contribute to the broad host recognition of this phage group^42,44^. In our HIDEN-SEQ library, several *bona fide* tail fiber genes (*bas54_0026, bas54_0027, bas54_0028, bas54_0031,* and *bas54_0041*) tolerated disruptive insertions, indicating that they are dispensable for infection of *E. coli* K-12 and for the formation of infective virions. We did not observe transposon insertions in the ~2 kb intergenic region (between *bas54_0134* and *bas54_0135*) that separates genes transcribed in opposite directions, suggesting an important functional role of this locus (Fig. 3b). Strikingly, 15 of 44 essential genes in Bas54 genome lack any functional annotation, highlighting that a substantial portion also of the essential genome remains elusive in phages beyond classical models. These results demonstrate that HIDEN-SEQ is both modular and flexibly applicable across different viral groups.

### HIDEN-SEQ reveals new anti-defense genes in different phages

The critical breakthrough of HIDEN-SEQ is its ability to assign new functions to the uncharacterized “dark matter” of phage genes by conditional essentiality in specific setups. After establishing the technology using the *E. coli* K-12 lab strain, we next asked whether HIDEN-SEQ could be applied also in a more complex and clinically relevant context. We thus used viral transposon libraries to infect a panel of *E. coli* clinical isolates from urinary tract infections obtained through the Swiss NCCR AntiResist consortium (Extended Data Table 1, *to be published elsewhere*) and compared gene essentiality for each phage between strains. To do so, we first selected a subset of clinical isolates based on sensitivity to the phages used in this study and infected them with transposon libraries of phages T4, Bas37, and Bas54. Intriguingly, for each phage-host pair, we identified multiple viral genes with significantly reduced insertion counts relative to the reference strain, indicating conditionally essential roles during infection of specific strains (Fig. 4a and Extended Data Fig. 6a, b). While these genes may contribute to diverse viral functions during infection, e.g., to exploit host-specific resources^45^, many are probably involved in overcoming bacterial immunity which is much more variable across *E. coli* strains than, e.g., metabolism^46^. In Bas54, some lateral tail fiber genes also showed host-specific conditional essentiality, suggesting that this phage deploys alternative tail fibers to target different host receptors (Extended Data Fig. 6c).

**Fig. 4.**
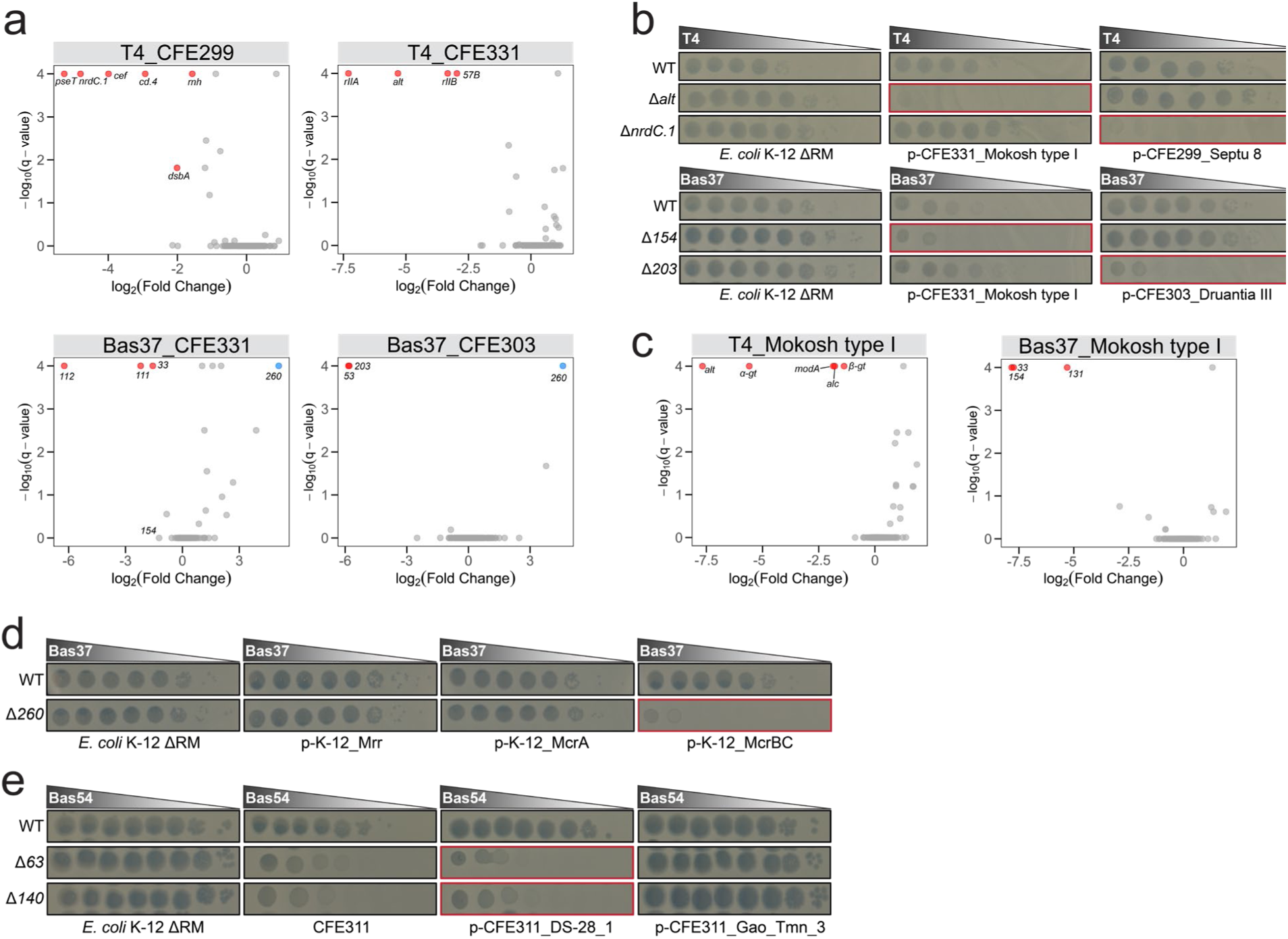
Identification of conditionally essential phage genes in clinical isolates by HIDEN-SEQ. **a**, Volcano plots comparing the depletion of gene disruptions in phages T4 and Bas37 after infection of laboratory strain *E. coli* K-12 BW25113 with infections of uropathogenic isolates *E. coli* CFE229, CFE331, and CFE303. Each point represents a viral gene with the *x*-axis showing the log₂(fold change) in normalized insertion counts and the *y*-axis showing the –log₁₀(*q*-value) (see *Methods*). Negative log₂(fold change) values indicate that mutants with insertions in that gene had reduced fitness in the clinical isolate compared to the laboratory strain. Genes highlighted in red represent the most significant observed effects. *bas37_0112* and *bas37_0111* encode RIIA and RIIB orthologs, respectively, and *bas37_0033* encodes for Alt. The single blue point represents disruptions in *bas37_0260* which causes a strong fitness defect in *E. coli* K-12 BW25113 but not in the clinical isolates. In Bas37_CFE331 plot, *bas37_0154* is also indicated because a phenotype with the knockout mutant was observed in the strain expressing the Mokosh type I of CFE331 isolate (see panel b). **b**, Ten-fold serial dilution plaque assays of T4 and Bas37 knockout mutants lacking candidate anti-defense genes plated on *E. coli* K-12 ΔRM variants expressing predicted defense systems from uropathogenic *E. coli* strains. Data are representative of 3 independent biological replicates. **c**, Volcano plots comparing the depletion of gene disruptions in phages T4 and Bas37 phage after infection of the *E. coli* K-12 ΔRM with infections of *E. coli* K-12 ΔRM variant expressing the Mokosh type I defense system of *E. coli* CFE331. Each point represents a viral gene with the *x*-axis showing the log₂(fold change) in normalized insertion counts and the *y*-axis showing the –log₁₀(*q*-value) (see *Methods*). Negative log₂(fold change) values indicate that mutants with insertions in that gene had reduced fitness in the Mokosh expressing strain compared to the control. *bas37_0131* encodes the ortholog of ADP-ribosyltransferase ModA. **d, e**, Ten-fold serial dilution plaque assays of Bas37 (**d**) or Bas54 (**e**) knockout mutants lacking candidate anti-defense genes plated on *E. coli* K-12 ΔRM variants expressing any of the three K-12 type IV restriction systems or predicted defense systems of *E. coli* CFE311. Data are representative of 3 independent biological replicates.

We first focused our follow-up experiments on the T-even phages which are known to harbor a diverse and effective repertoire of anti-defense factors^47,48^. Notably, we observed overlapping sets of conditionally essential genes for the T4 and Bas37 libraries grown on the same clinical strains (Fig. 4a and Extended Data Fig. 6a). However, additional factors became essential when Bas37 was grown on strains that T4 could not infect. To link these candidate anti-defense genes to specific branches of bacterial immunity, we used DefenseFinder^49^ to predict defense genes in these hosts and selected some defense systems for further analysis. In parallel, we generated targeted deletions of a subset of the most strongly depleted phage genes (log₂FC < −2). We then tested the growth of each phage mutant in the presence of each cloned defense system to identify possible phage gene / host defense pairings (Extended Data Fig. 7a, b). Remarkably, this screen revealed multiple phage anti-defense genes acting against different defense systems (Fig. 4b), including both novel factors and genes with known functions.

One of these cases was the discovery that Alt of T4 phage and β-α glucosyltransferase (*βα-gt*; *bas37_0154*) of Bas37 are involved in evasion of the Mokosh type I defense system of *E. coli* strain CFE331 (Fig. 4b). The *alt* gene encodes an ADP-ribosyltransferase that has been previously reported to manipulate the host RNA polymerase via ADP-ribosylation^50^ and to interfere with the MazEF toxin-antitoxin system^51^. We compared the Alt variants of both phages which exhibit notable sequence differences (Extended Data Fig. 8a) and tested whether they could function interchangeably. When expressed ectopically in the presence of Mokosh type I, Alt^T4^, but not Alt^Bas^^37^ (Bas37_0033), partially restored plaque formation of the T4 *alt* mutant, Bas37 mutants, and also Bas54, which is otherwise sensitive to Mokosh (Extended Data Fig. 8b). Together with the sensitivity of the Bas37 *βα-gt* (*bas37_0154*) knockout mutant, these results indicated that complete evasion of Mokosh type I depends on the glucosylation of viral DNA and the ADP-ribosylation of host proteins which was confirmed using T4 and Bas37 HIDEN-SEQ libraries (Fig. 4c; see also *Supplementary Discussion*).

We also identified *nrdC.1*, a T4 gene of unknown function, as *a bona fide* inhibitor of the Septu defense system of *E. coli* strain CFE299. The protein encoded by this gene has no known functional domains and no significant structural similarity to previously characterized proteins (Extended Data Fig. 8c). Similarly, the *bas37_0203* gene, also of unknown function, was identified as a *bona fide* inhibitor of Druantia type III of *E. coli* strain CFE303. Like *nrdC.1*, this gene lacks any known domains or characterized homologs (Extended Data Fig. 8d), so the mechanism of action remains elusive for both *bona fide* anti-defense factors. One additional interesting finding relates to Bas37 gene *bas37_0260* which is encoded at the same locus as the T4 inhibitor of the GrmSD type IV restriction system called internal protein I (IPI)^52^ (Extended Data Fig. 8e). Disruption of *bas37_0260* caused a strong fitness defect when infecting the *E. coli* K-12 BW25113 laboratory strain but not the different uropathogenic strains or the closely related *E. coli* K-12 MG1655 ΔRM variant in which we had deleted the three type IV restriction systems McrA, McrBC, and Mrr (Fig. 4a and Extended Data Fig. 6a; see also Extended Data Table 7)^42^. To confirm that Bas37_0260 acts as an inhibitor of *E. coli* K-12 type IV restriction systems, we deleted this gene from Bas37 phage and found that the knockout mutant became specifically sensitive to McrBC (Fig. 4d). These data demonstrate that *bas37_0260* gene encodes a previously unknown type IV restriction inhibitor which shares no homology to the known analogs IPI, IPII, and IPIII of phage T4 other than the characteristic N-terminal signal sequence that directs its packaging into the capsid (Extended Data Fig. 8f)^53^.

Lastly, we extended the phage gene / host defense pairing screen to Bas54 phage and identified two anti-defense genes specifically required to overcome DS-28, a predicted defense system encoded by *E. coli* strain CFE311 (Fig. 4e). *bas54_0063* gene encodes for a MoxR ATPase while *bas54_0140* is a gene of unknown function.

Together, these findings show that HIDEN-SEQ can systematically uncover phage genes that counteract bacterial defenses, including those with no prior annotations or characterized function.

## Discussion

In this study, we have established HIDEN-SEQ as the first genome-wide transposon-insertion sequencing method for bacteriophages. This method allows systematic disruption of phage genes by Himar1 transposition and selection for transposon mutants with a CRISPR-Cas13a / anti-CRISPR system. Using HIDEN-SEQ, we constructed highly saturated transposon libraries for three different phages that allowed us to systematically map phage gene essentiality at unprecedented resolution.

We benchmarked the technique using the well-studied T4 model phage, where most genes are dispensable under standard laboratory conditions^10^, and show that HIDEN-SEQ reliably recapitulates its known biology. Only a few genes (*inh, hoc, segE, uvsW, segC,* and *wac*) appeared essential in our screen despite being reported as not essential in earlier studies^10^. The high saturation of our library before selection (Extended Data Fig. 2a) confirms that transposition is not the limiting factor and that post-selection depletion reflects real fitness effects of insertions in our experimental setup. One possible explanation for these few observed differences to previous work may be the different strategies used to generate viral mutants. For example, in our dataset the essentiality of *wac*, encoding the phage whiskers, is compared to earlier work which had studied conditional translational inhibition by amber mutants^54^, which may allow residual protein expression. Although transposon insertions can in some cases affect nearby genes^24^, our data argue against widespread polar effects because insertions upstream of essential genes were usually well tolerated (see Figs. 1b and 3b). Nevertheless, locus-dependent effects such as altered downstream gene expression remain possible and may explain some of these discrepancies.

Beyond T-even phages, HIDEN-SEQ revealed many genes of unknown function with essential roles in non-model phage infection, highlighting its potential to explore gene function across diverse phage groups. A major strength of this method is its ability to uncover conditionally essential genes that become critical only under certain challenges. HIDEN-SEQ not only identified known anti-defense genes in the presence of their corresponding bacterial defense systems but also uncovered new *bona fide* factors acting against six distinct defense systems (DarTG2, Mokosh type I, Septu, Druantia type III, McrBC type IV restriction, and DS-28). Five of these viral genes had no previously known function, demonstrating the unique ability of HIDEN-SEQ screens to gain access and assign functions to the “dark matter” of phage genomes.

Several approaches for mapping phage gene essentiality have recently expanded the toolbox for phage biology and feature specific disadvantages compared to HIDEN-SEQ^18–23^. The knockdown-based method CRISPRi-ART^19^, for instance, relies on previous knowledge about the position of viral genes and accurate start codon prediction while also being constrained by the notorious variability of crRNA efficacy. Consequently, this method failed to detect the essentiality of around 25% of the known essential genes of phage T4. Moreover, CRISPRi-ART relies on knockdown efficacy in all tested conditions, which further restricts the scalability of this method compared to robust gene disruptions by transposition in HIDEN-SEQ. However, knockdown approaches such as CRISPRi^18,19^ or ASO-based methods^23^ are suited to probe fitness effects of essential genes which are inherently inaccessible to HIDEN-SEQ. Another recently published method, PhageMaP, demonstrated that phage gene essentiality can be mapped by targeted mutagenesis^20^. However, this is a laborious approach that requires extensive custom construct design and cloning for each phage. In addition, the method generates only a few insertional mutants per gene, chosen in a targeted manner, which introduces bias and limits resolution. In contrast, HIDEN-SEQ requires no prior specific knowledge about genes in the target phage, relies on a single crRNA for selection per phage, and generates high-density insertion libraries that can be repeatedly used to study phage infection across different hosts and conditions. Compared to existing approaches, HIDEN-SEQ thus achieves a unique combination of experimental utility and high resolution, enabling systematic mapping of dispensable and conditionally essential phage genes.

Taken together, HIDEN-SEQ provides a versatile platform that links phage phenotypes to underlying genes and, consequently, molecular mechanisms. It enables high-throughput mapping of phage-host interactions beyond standard laboratory conditions, including the infection of clinical isolates and under varied growth environments. In addition, HIDEN-SEQ reveals dispensable genome regions that can be directly informative in phage engineering for clinical and biotechnological applications. Looking forward, its unique utility and efficacy make HIDEN-SEQ a powerful tool to address increasingly complex biological questions about phage biology with far-reaching implications for phage therapy, microbial ecology, and biotechnological innovation.

## Supporting information

Escherichia virus T4D genome

## Methods

### Preparation of culture media and solutions

Lysogeny Broth (LB) was prepared by dissolving 10 g/l tryptone, 5 g/l yeast extract, and 10 g/l sodium chloride in Milli-Q H_2_O, then sterilized by autoclaving. LB agar plates were prepared by supplementing LB medium with 1.5% w/v agar prior to autoclaving. M9 minimal medium supplemented with 0.4% w/v D-glucose (M9Glc) was prepared as previously described^55^. M9Glc agar plates were prepared by mixing autoclaved 3% (w/v) agar and sterile 2× M9Glc medium in a 1:1 ratio to achieve a final agar concentration of 1.5%. Phosphate-buffered saline (PBS) was prepared as a solution containing 8 g/l NaCl, 0.2 g/l KCl, 1.44 g/l NA_2_HPO_4_·2H_2_O, and 0.24 g/l KH_2_PO_4_, adjusted to pH 7.4 with 10 M NaOH and sterilized by autoclaving. SM buffer was prepared as 0.1 M NaCl, 10 mM MgSO_4_, and 0.05 M Tris (pH 7.5) and was also sterilized by autoclaving.

### Bacterial strains handling and culturing

*E. coli* strains were routinely cultured in LB medium at 37°C with agitation at 170 rpm. For experiments involving the DarTG2 system, the bacteria were instead cultured in M9Glc medium and incubated at 30°C^56^. We selected for plasmid maintenance using ampicillin at 50 µg/ml, chloramphenicol at 25 µg/ml, gentamicin at 20 µg/ml, and kanamycin at 25 µg/ml. A full list of bacterial strains used in this study is provided in Extended Data Table 1.

### Bacteriophage handling and culturing

Bacteriophages were cultured using the double-agar overlay (“top agar”) method^57^. Top agar was prepared as LB agar with 0.5% w/v agar and supplemented with 20 mM MgSO_4_ and 5 mM CaCl_2_. Top agar plates were incubated at 37°C or 30°C (Δ*adfM* T4 phage experiments) for ca. 16-20 h before plaque enumeration. For CRISPR-Cas13a counterselection, top agar was made without divalent cations and supplemented with chloramphenicol at 25 µg/ml and anhydrotetracycline (aTc) at 5nM to induce the expression of Cas13a^33^. For transposition experiments, 2 mM IPTG (Isopropyl-β-D-thiogalactopyranoside) was added to the top agar to induce transposase expression. When M9Glc plates were used, top agar was prepared by mixing sterile 1% w/v agar with 2× M9Glc medium in a 1:1 ratio, to have a final agar concentration of 0.5% w/v and maintain equivalent nutrient composition to the M9Glc agar plate. All bacteriophages used in this study are listed in Extended Data Table 2.

### Plasmids and plasmid construction

Plasmids were typically constructed using the method of Gibson et al.^58^ (“Gibson Assembly”) by ligating PCR products containing 25 nucleotide overlaps. Some plasmids have been constructed using classical restriction-ligation cloning as described in our previous work^42^. Mutations in plasmids were introduced by round-the-horn PCR^59^ with partially overlapping primers. We used a two-plasmid setup for LbuCas13a counterselection, where we separated Cas13a protein expression into a dedicated plasmid pAH221 (derivative of all-in-one plasmid pBA559^33^; Addgene 186235) and cloned the crRNA expression cassette into pAH218 (RSF1010 origin of replication). The same two-plasmid setup was used to establish the LseCas13a system in *E. coli*. The coding sequence of the LseCas13a protein was codon-optimized for *E. coli* and synthesized by Twist Bioscience before cloning into the pAH221 backbone. In parallel, the crRNA expression cassette was cloned into the pAH218 backbone. The crRNAs for selection against wildtype phages were designed to target essential phage transcripts, using either 31 (LbuCas13a) or 24 nucleotide (LseCas13a) spacers^31,33^ and cloned into the pAH218 backbone via Golden Gate assembly^33^. Codon-optimized gene sequences for the anti-CRISPRs, AIcrVIA3 (LbuCas13a)^43^ and AcrVIA1 (LseCas13a)^30^, were synthesized (Twist Bioscience) and cloned into the transposon and homologous recombination vectors. Transposon plasmids (RSF1030 high-copy origin of replication) contained *mariner* transposon inverted repeats (GGGGACTTATCAGCCAACCTGTTATGT) flanking the selection marker. The transposase gene was cloned into pAH186, a ColE1-based plasmid with IPTG-inducible expression. Defense systems were predicted in the genome of the *E. coli* clinical isolates using DefenseFinder (v2.0.0)^49^ (Extended Data Table 3, *detailed predictions will be available upon publication*) and then cloned on a SC101-*ori* plasmid under their native promoter (by including >80 bp upstream region) and used to transform *E*. *coli* K-12 MG1655 ΔRM^42^ for further testing. For phage gene complementation, the pNDM220 backbone was used, which contains an IPTG-inducible P*lac* promoter on a mini-R1 backbone. All cloned constructs were routinely verified by Sanger Sequencing or full plasmid sequencing using Oxford Nanopore technology at Microsynth AG. The construction of all plasmids is described in detail in Extended Data Table 4. All oligonucleotide primers and synthesized DNA fragments used in this work are listed in Extended Data Table 5.

### Construction of phage gene knockouts

Knockout mutants of bacteriophage genes were constructed by replacing the coding sequence of the target gene with an *acr* gene through homologous recombination followed by CRISPR-Cas selection as pioneered by Guan and colleagues^31^. Briefly, the *acr* gene was cloned with flanking >100 bp phage homology arms onto a replicative plasmid as recombination donor. Subsequently, an *E. coli* K-12 host was transformed with this plasmid and infected with the target phage to enable homologous recombination. The lysate from this infection was then subjected to CRISPR-Cas13a selection against the parental wildtype phage using a crRNA targeting the transcripts of an essential phage gene. Subsequently, plaques were screened for successful insertion of the *acr* gene by overspanning PCR and the sequence of the recombinant phage was verified by Sanger Sequencing at the relevant locus. The phage mutant isolates were passaged three times via single plaques on top agar of the Cas13a selection strain for clonal purification.

### Construction of the T4 HIDEN-SEQ library

The transposition donor strain was obtained by sequentially transforming *E. coli* K-12 with two plasmids. The first plasmid encoded the Himar1 transposase under the control of an IPTG-inducible promoter, while the second carried the *mariner* transposon *in trans* on a high-copy plasmid to bypass the notorious overexpression inhibition of *mariner*-type transposases^60^. For the T4 HIDEN-SEQ library, the transposon encoded the *acrVIA1* gene without a transcriptional terminator to allow readthrough transcription. It also included stop codons in all three reading frames at the 3’ end of the *acrVIA1* gene to prevent aberrant translation. To generate a transposon pool, phage T4 was grown to confluency on top agar plates of the transposon donor strain supplemented with IPTG for transposase induction. Confluent plaques from these plates were scraped and pooled in 10-15 mL of SM buffer, then centrifuged at 8000 × *g* for 10 min. Supernatants were collected and titers were determined, typically yielding >10^10^ PFU/ml. Even at possibly very low transposition frequencies like 1 in 10^5^ clones, at least ≥10^5^ PFU/mL would be transposon mutants, which is well above the ~15,000 unique insertions needed to potentially saturate all 14,474 TA sites in the genome of our T4 lab variant (Escherichia_virus_T4D.fasta, provided as Supplementary File; see also Extended Data Table 2). This lysate was then plated on top agar with the selection strain expressing LseCas13a and a crRNA targeting the transcripts of the major capsid protein (*mcp*). As a control, a parallel lysate was prepared using a donor strain carrying a non-selectable transposon without anti-CRISPR and plated on the same selection strain to estimate the frequency of Cas13a genetic escape. We tested different volumes of the phage lysate for plating to obtain single plaques with adequate spacing where direct competition between mutants may be reduced. Plaques were counted to estimate the number of single plaques per square plate, which was used to calculate the number of plates required to make the library. We aimed to collect at least twice as many plaques as there are TA dinucleotide sites in the phage genome to ensure high saturation. Plaques from all plates were scraped and pooled in 100-200 mL of SM buffer and centrifuged at 8000 × *g* for 10 min. The resulting final stock of the library had a titer of 10^9^-10^10^ PFU/ml, predominantly transposon mutants with a possible minor fraction of genetic escape mutants. The library was stored at 4°C till further use. Following the same protocol, a second T4 library was later constructed on the *E. coli* Δ*rnlA* strain from the Keio collection^61^, which does not express the RnlA-RnlB system, to test the conditional essentiality of *dmd* (Fig. 2b).

### Construction of Bas37 HIDEN-SEQ library

The Bas37 HIDEN-SEQ reference library was constructed using the same procedure as for T4. For counterselection, we used the same crRNA targeting the major capsid protein, which was also effective against Bas37 because the target sequence is conserved between these two closely related phages^42^. We collected a similar number of plaques aiming to saturate the 14,279 TA dinucleotide sites in the Bas37 genome^42^.

### Construction of Bas54 HIDEN-SEQ library

The Bas54 HIDEN-SEQ reference library was generated using the same procedure described above with a simple adaptation of the selection system to this virus. For Bas54, the transposon encoded the anti-CRISPR gene *AIcrVIA3*, and selection was consequently performed using the LbuCas13a system with crRNA targeting the transcripts of the DNA polymerase gene. For the first library, we estimated that ~17,000 plaques were pooled to cover 8,585 TA dinucleotide sites in the Bas54 genome^42^, following our general library construction approach. In the second library construction attempt, we collected roughly twice as many plaques as in the first to ensure higher saturation of the transposon library.

### Competitive fitness assay using HIDEN-SEQ library

Overnight cultures of the target hosts were diluted 1:10 to mid-exponential phase (ca. 5×10^8^ CFU/ml) in fresh LB medium with additional supplements if necessary. The HIDEN-SEQ reference phage library was added at an MOI of 1:100 to ensure single infections. At this MOI, infections are performed with 5×10^6^ PFU/ml which ensures that there are more phages per milliliter than the transposon mutant pool can contain independent mutants. This excess robustly buffers the experiment against random perturbations of mutant abundance. After agitation at 170 rpm in Erlenmeyer flasks for 20-24 h at 37°C, the cultures were centrifuged at 8000 × *g* for 10 min, and the supernatants were transferred to fresh tubes. Mutant library phage lysates were stored at 4°C prior to genomic DNA extraction.

Condition-specific modifications were applied when necessary: For CBASS experiments, 0.2% L-arabinose was added at the time of infection to induce expression of defense genes^37^. Competitive fitness assays to test growth medium-dependent conditional essentiality were performed in both LB and M9Glc medium. For experiments with the DarTG2 system, infections were performed in M9Glc medium with incubation at 30°C^56^. Further details of all libraries generated in this work are provided in Extended Data Table 6-8.

### HIDEN-SEQ library preparation and sequencing

Genomic DNA was extracted from phage transposon mutant library samples using Norgen Biotek Phage DNA Isolation Kit. Library preparation and sequencing were performed at the Lausanne Genomic Technologies Facility (GTF) using the AVITI short-read sequencer (Element Biosciences), following a customized protocol. Briefly, genomic DNA was sheared to ~300-500 bp fragments and ligated to Illumina-compatible adaptors containing i5 and i7 indexes. A first round of PCR was performed using a transposon-specific primer with a 5’ biotin label and an adaptor-specific primer to selectively amplify transposon insertion sites. Biotinylated PCR products were captured using streptavidin-coated magnetic beads to remove background genomic DNA. Finally, to enrich transposon-insertion sites, another PCR was performed using a second transposon-specific primer that anneals close to the transposon-genome junction and incorporates the P5 flow cell adapter. The resulting libraries were deep sequenced using single-end sequencing.

### Data analysis of HIDEN-SEQ libraries

Raw sequencing reads were initially assessed for quality using FastQC (v0.12.1). Adapter trimming and quality filtering were subsequently performed with fastp (v0.22.0)^62^, applying a minimum Phred quality score cutoff of Q30 and discarding reads shorter than 50 bp. The preprocessed reads were then used to identify transposon insertion sites with the TPP tool from TRANSIT (v3.3.13) software^63^. TPP first detects a match (allowing no mismatch) to the transposon prefix in each read and then maps the genomic portion to the phage reference genome using BWA (v0.7.12)^64^. The output of this step is a read count table of transposon insertions at every TA dinucleotide site in the genome. To reduce noise from spurious reads, TA sites with 10 or fewer mapped reads were set to zero. Due to the variable density of TA sites across different phages, we implemented a random read subsampling strategy using seqkit (v2.10.0)^65^ to standardize the analysis across different datasets. Specifically, for each phage, we estimated a target sequencing depth of 350 reads per TA site, which maintains the library complexity and ensures that the global sequencing depth is normalized across genomes with different insertion site availability.

### Gene essentiality analysis

To account for the less complex structure and smaller size of phage genomes compared to bacterial genomes^29,63^, we developed a custom gene essentiality classification pipeline in RStudio (v4.3.2). This approach integrates insertion site saturation, positional bias, and read counts to more accurately capture the range of fitness effects observed in our data, which did not conform solely to binary essentiality classifications. For each gene, we calculated saturation as the proportion of TA sites with non-zero read counts. Genes with saturation below 0.2 were classified as essential, while those with saturation above 0.8 were classified as non-essential. Genes falling between these thresholds were further evaluated in two steps. We first assessed the positional distribution of insertions. Genes with a strong 3′ end insertion bias or with insertions restricted to one half of the gene, indicative of domain-level essentiality (if genes encode proteins ≥80 aa), were also classified as essential. We then used read count as a proxy for the fitness cost of gene disruptions. Genes with saturation between 0.2–0.4 were classified as “strong fitness defect” if more than 50% of insertions had read counts below the dataset median, indicating that gene disruption imposes a high fitness cost. In contrast, genes with saturation between 0.7–0.8 were classified as “reduced fitness” if the majority of insertions had read counts above the median, indicating a non-essential gene whose disruption has a mild fitness cost. Genes with saturation between 0.4 and 0.7, or those that did not meet the above criteria, were classified as “intermediate”, reflecting phenotypes that were neither clearly deleterious nor fully tolerated. Each gene was thus ultimately assigned to one of five HIDEN-SEQ categories: essential, strong fitness defect, intermediate, reduced fitness, or non-essential. Genes containing ≤ 5 TA sites were omitted from classification due to insufficient insertion sites to support confidence calls. To account for stochastic variation introduced by subsampling, we generated five replicate datasets by randomly sampling from the same set of preprocessed reads at fixed sequencing depth. Each replicate was processed independently through the full analysis pipeline. Final gene essentiality calls were assigned based on consistent classification in at least three out of five replicates. Insertions at TA sites outside of coding sequences were recorded and analyzed as described above but not subjected to an essentiality analysis.

### Conditional essentiality analysis

To identify genes that become essential only under specific experimental conditions, we used the resampling method implemented in TRANSIT^63^, which determines read counts that are significantly different between two conditions using a non-parametric permutation test. To account for differences in sequencing depth between datasets, we used the subsampling at a fixed depth, as described above, and normalized the read counts using the Trimmed Total Reads (TTR) method^63^. Instead of comparing each condition to the original reference library, we used parallel control libraries that were grown under the same conditions but lacking selective pressure. As an example, infection of a host carrying an empty vector was used as a control to infections of the same host carrying a plasmid expressing an anti-phage defense system. This approach minimized noise from potential random loss of mutants during library passaging and helped distinguish true conditionally essential genes from those that get depleted after prolonged selection (see *Supplementary Discussion*). The resampling test was run with 10,000 permutations (-s 10000), and returned, for each gene, a *P*-value and the corresponding log_2_ fold-change (log_2_FC) in normalized insertion counts between conditions. The *P*-values were adjusted for multiple testing by the Benjamini-Hochberg procedure to control the false-discovery rate at < 0.05 (*q*-value). Given the small size of the phage genomes, transposon insertion profiles were also inspected manually as a complementary check. For the CBASS condition, resampling results were not considered due to random mutant losses on this sample (because of toxicity of defense genes), however manual inspection revealed one gene with clear gene-wide depletion. In the Bas54 HIDEN-SEQ library grown in *E. coli* CFE303, the *bas54_0108* gene showed clear depletion relative to the reference strain but was not detected by the standard procedure due to high insertion counts at the 3′ end. To account for this, we applied trimming of TA sites in the C-terminus (-iC 20) during resampling.

### Genome sequencing and assembly of individual phage mutants

Whole-genome sequencing of individual phage mutants was performed using commercial DNA sequencing as described previously^42,66^. Briefly, phage genomic DNA was isolated from the lysate using the Norgen Biotek Phage DNA Isolation Kit and sequenced at SeqCenter (https://www.seqcenter.com/) with Illumina technology. Genome assembly was done using Unicycler (v0.4.8)^67^ and the downstream analyses were performed on Geneious Prime 2023.0.4.

### Structure predictions and analysis of newly identified anti-defense genes

To generate hypotheses about the function of the candidate anti-defense phage genes, we searched for structural homologs in the Protein Data Bank using HHpred^68^ and Dali server^69^. Structure predictions of phage proteins were performed using AlphaFold2 on the ColabFold webserver^70^ with default parameters. The structural visualizations were generated using PyMOL Molecular Graphics System (Version 3.0.4, Schrodinger LLC).

## Acknowledgments

The authors thank Dr. Michele LeRoux for generously sharing the DarTG plasmids and Prof. Philip J. Kranzusch for the CBASS plasmids. We thank Prof. Urs Jenal for sharing the *E. coli* K-12 Δ*rnlA* BW25113 Keio collection mutant. The clinical isolates from the collection of Swiss NCCR AntiResist consortium were obtained from Prof. Christoph Dehio. We thank Dr. Enea Maffei and Kamile Cerepenkaite for help with plasmids. Dr. Afonso M. Bravo and Dr. Jesús Cámara-Almirón are acknowledged for advice regarding the sequencing of transposon insertion sites. The authors thank Prof. Melanie Blokesch for advice and suggestions. We thank the Lausanne Genomic Technologies Facility (GTF) for efficient, knowledgeable, and reliable sequencing of HIDEN-SEQ libraries. This work was supported by the Swiss National Science Foundation (SNSF) Ambizione Fellowship PZ00P3_180085 (to A.H.), SNSF Starting Grant TMSGI3_211369 (to A.H.), and SNSF National Centre of Competence in Research (NCCR) AntiResist (grant number 180541).

## Author Contributions

A.H. and D.H. developed HIDEN-SEQ with advice of J.W.V. throughout the project. Initial experiments have been performed by A.H., while the final setup of HIDEN-SEQ and its application to T4, Bas37, and Bas54 was implemented by D.H., with help of J.R. Cloning and analysis of defense systems in uropathogenic *E. coli* strains have been performed by D.P. D.H. and A.H. wrote the manuscript with input from all authors.

## Additional Information

Supplementary Information is available for this paper. Correspondence and requests for materials should be addressed to Prof. Alexander Harms.

## Data Availability

Source data of the experiments in this study are included as data files with this study.

## Code Availability

The R-studio code used for the gene essentiality analysis of HIDEN-SEQ libraries is available at Zenodo with the digital object identifier (*to be released*).

## Extended Data Figures and Tables

**Extended Data Fig. 1.**
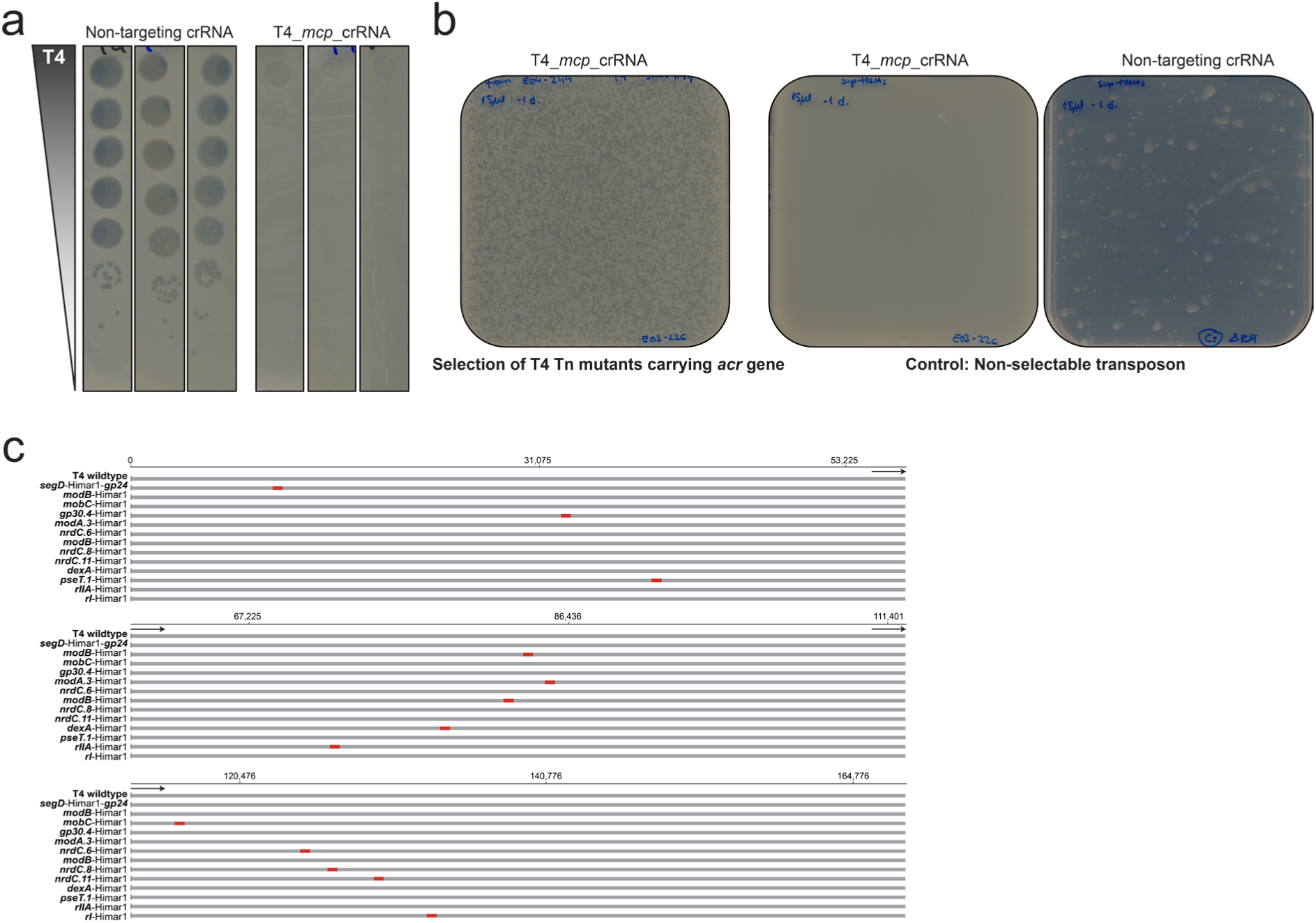
The CRISPR-Cas13a / anti-CRISPR based selection of transposition events into T4 phage genome. **a**, Ten-fold serial dilution plaque assays of T4 phage plated on *E. coli* K-12 ΔRM expressing *cas13a* with either a non-targeting crRNA control or a crRNA targeting the transcripts of the major capsid protein. **b**, T4 pool of anti-CRISPR transposon mutants plated on *E. coli* K-12 expressing *cas13a* with the crRNA targeting the transcripts of the major capsid protein. The plaques indicate phage growth enabled by anti-CRISPR activity (left). A T4 mutant pool generated with a non-selectable transposon (lacking anti-CRISPR) plated on the same host does not show plaque growth in presence of CRISPR-Cas13a selection (middle plate) but results in full lysis (right) without selection. **c**, Whole-genome alignment of wildtype T4 phage with 13 individual clones that were randomly picked after CRISPR-Cas13a selection and then sequenced. Red marks indicate transposon insertions identified in each clone. Alignments were generated using MAFFT v7.490 implemented in Geneious Prime 2023.0.4 with default settings.

**Extended Data Fig. 2.**
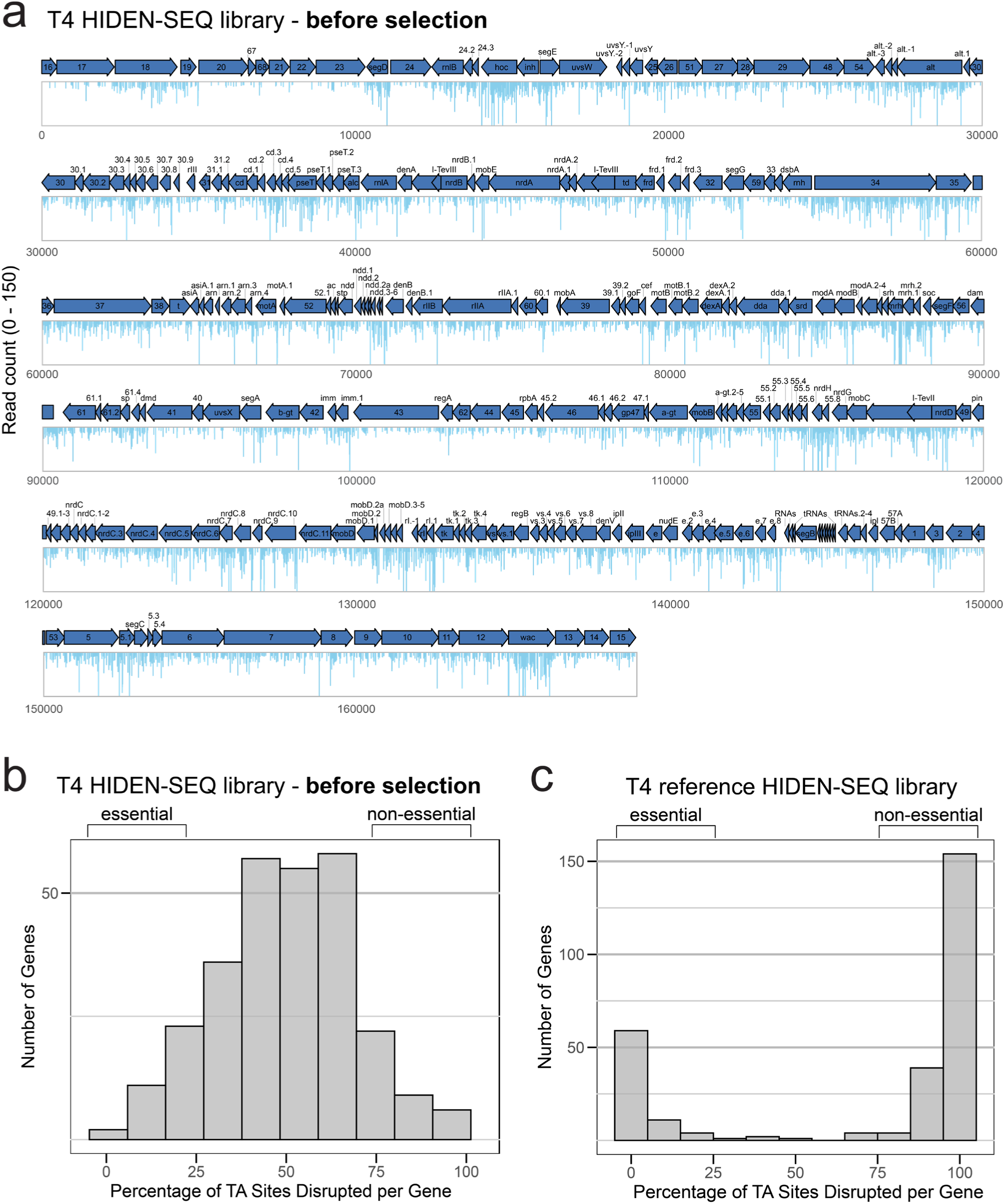
Genome-wide essentiality mapping of T4 phage. **a**, Transposon insertion profiles of the T4 input HIDEN-SEQ library collected directly after the transposition step, i.e., before significant selection against non-viable insertion mutants by viral replication. The start of the genome map has been set to the start of the small terminase subunit gene. Arrows present the coding sequences with the orientation indicating the direction of transcription. Only the full-length gene products are shown. Vertical bars represent the number of transposon insertions at TA dinucleotide sites with heights indicating read counts (up to a maximum of 150). **b**, **c**, Histograms illustrating the percentage of disrupted TA sites per gene for the T4 input HIDEN-SEQ library dataset (**b**) or the T4 reference HIDEN-SEQ library (**c**). The comparison shows that non-essential and essential genes are easily distinguishable in a highly saturated library.

**Extended Data Fig. 3.**
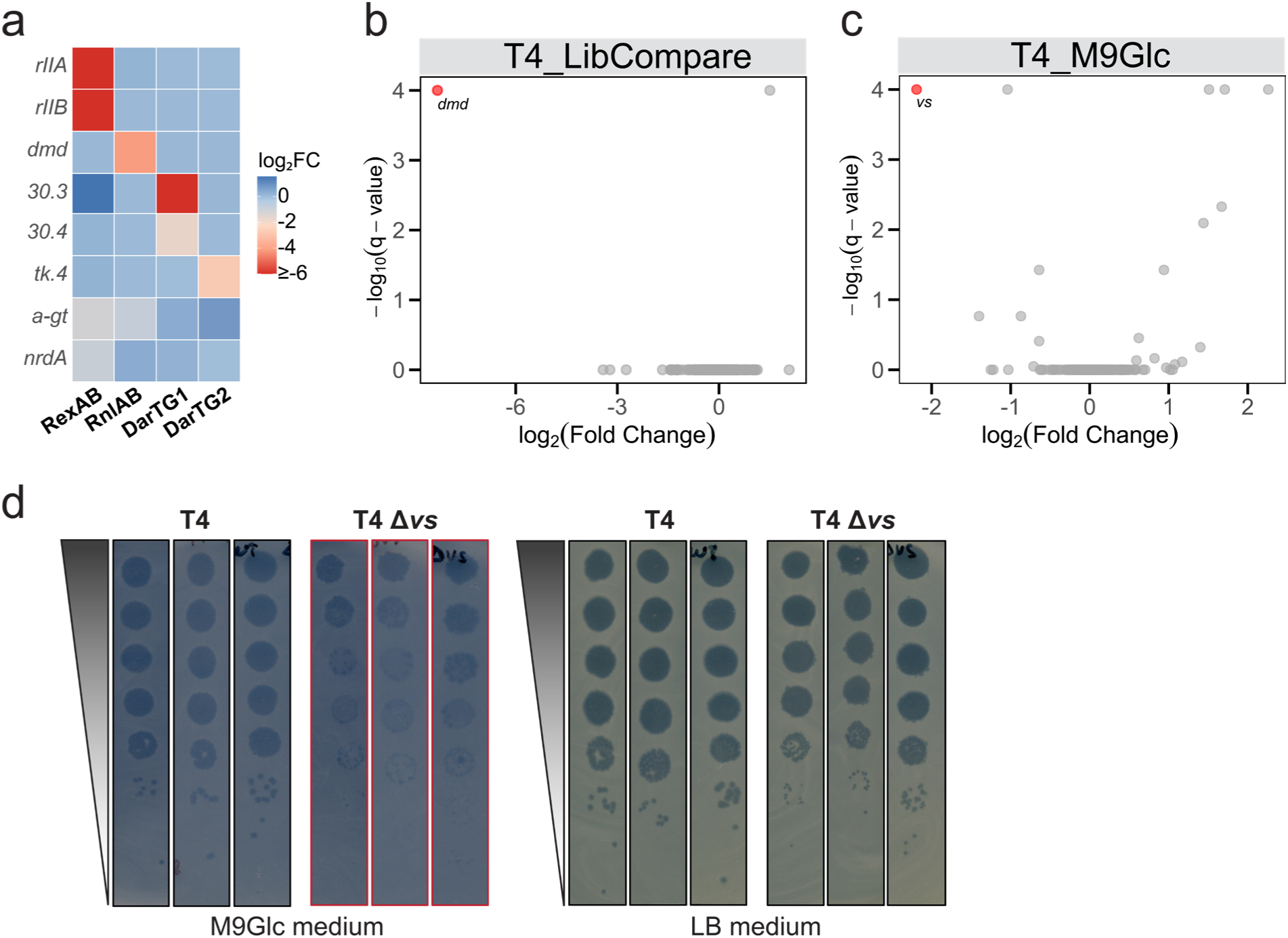
Context-dependent essentiality of T4 phage genes. **a**, Summary heatmap of conditionally essential genes identified by HIDEN-SEQ (see also figS. 2a-e). Genes with significant depletion determined by resampling analysis for all conditions are shown. **b**, Volcano plot comparing the depletion of gene disruptions in phage T4 between the reference HIDEN-SEQ library and the library generated on the *E. coli* K-12 *rnlA* knockout. Each point represents a viral gene with the *x*-axis showing the log₂(fold change) in normalized insertion counts and the *y*-axis showing the –log₁₀(*q*-value) (see *Methods*). **c**, Volcano plot comparing the depletion of gene disruptions in phage T4 after infection of *E. coli* K-12 BW25113 laboratory strain in rich LB medium and in M9Glc minimal medium. Each point represents a viral gene with the *x*-axis showing the log₂(fold change) in normalized insertion counts and the *y*-axis showing the –log₁₀(*q*-value) (see *Methods*). Negative log₂(fold change) values indicate that mutants with insertions in that gene had reduced fitness in the M9Glc medium condition compared to the LB medium. **d**, Ten-fold serial dilution plaque assays of wildtype T4 and a *vs* knockout mutant plated on *E. coli* K-12 BW25113 for growth on agar plates prepared with M9Glc or LB medium. The reduced growth of a viral *vs* mutant in the M9Glc minimal medium is apparent.

**Extended Data Fig. 4.**
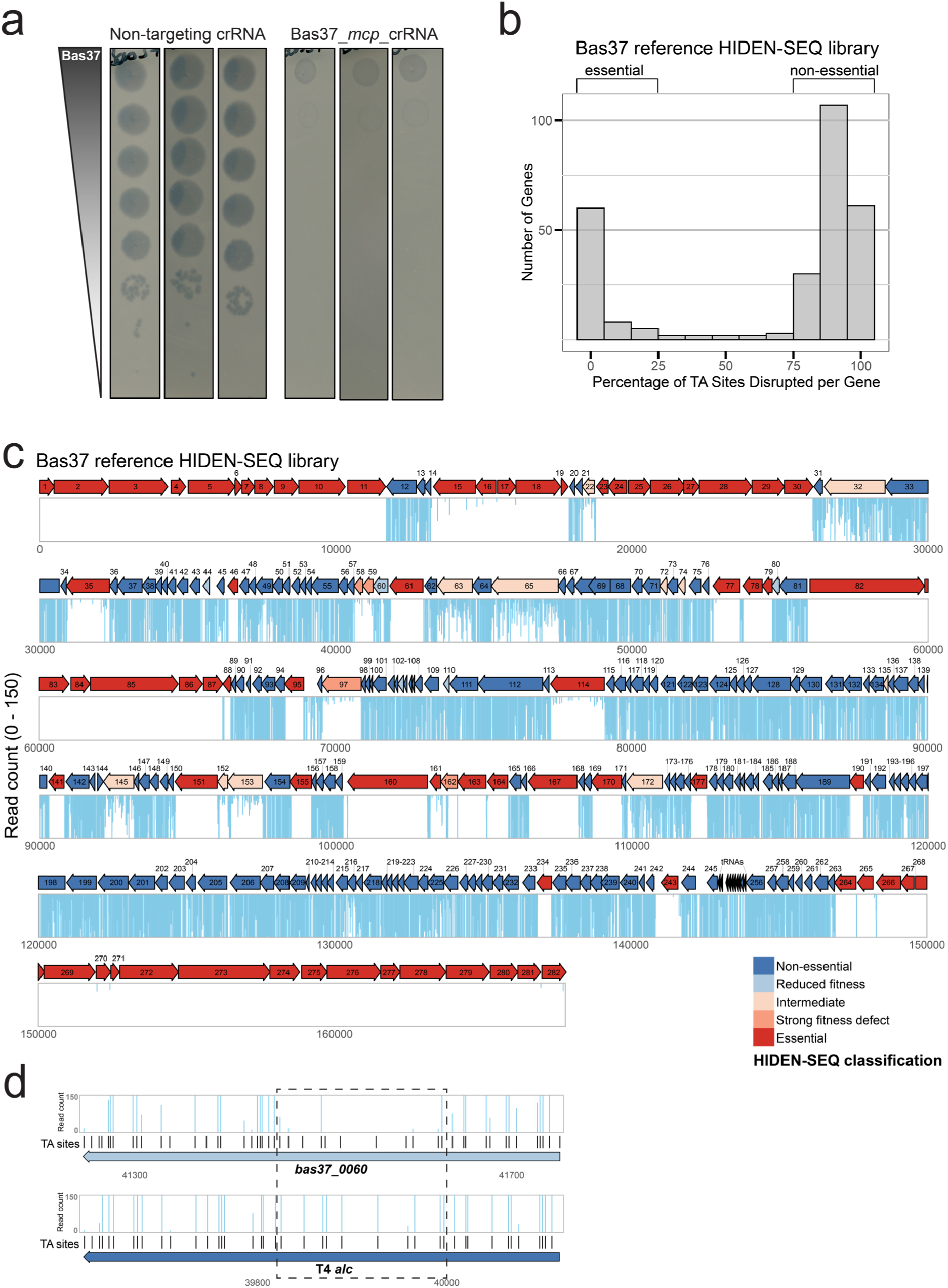
Genome-wide essentiality mapping of Bas37 phage. **a**, Ten-fold serial dilution plaque assays of phage Bas37 plated on *E. coli* K-12 ΔRM expressing *cas13a* with either a non-targeting crRNA control or a crRNA targeting the transcripts of the major capsid protein. **b**, The histogram illustrates the percentage of disrupted TA sites per gene for the Bas37 reference HIDEN-SEQ library showing that non-essential and essential genes are easily distinguishable in a highly saturated library. **c**, Phage Bas37 HIDEN-SEQ gene essentiality map with position 1 set to the start of the small terminase subunit gene. Arrows present the coding sequences with the orientation indicating the direction of transcription. Arrow colours correspond to HIDEN-SEQ classification categories derived from transposon insertion profiles: non-essential (dark blue), reduced fitness (light blue), intermediate (light orange), strong fitness defect (orange), and essential (red). Vertical bars represent the number of transposon insertions at TA dinucleotide sites with heights indicating read counts (up to a maximum of 150). Genes containing ≤5 TA sites are shown in black and were not classified. **d**, Transposon insertion profiles of *bas37_0060* (as an example) and its T4 ortholog *alc* gene highlighting that saturation differences (0.76 and 0.97) shifted the classification to reduced fitness in Bas37 and non-essential in T4.

**Extended Data Fig. 5.**
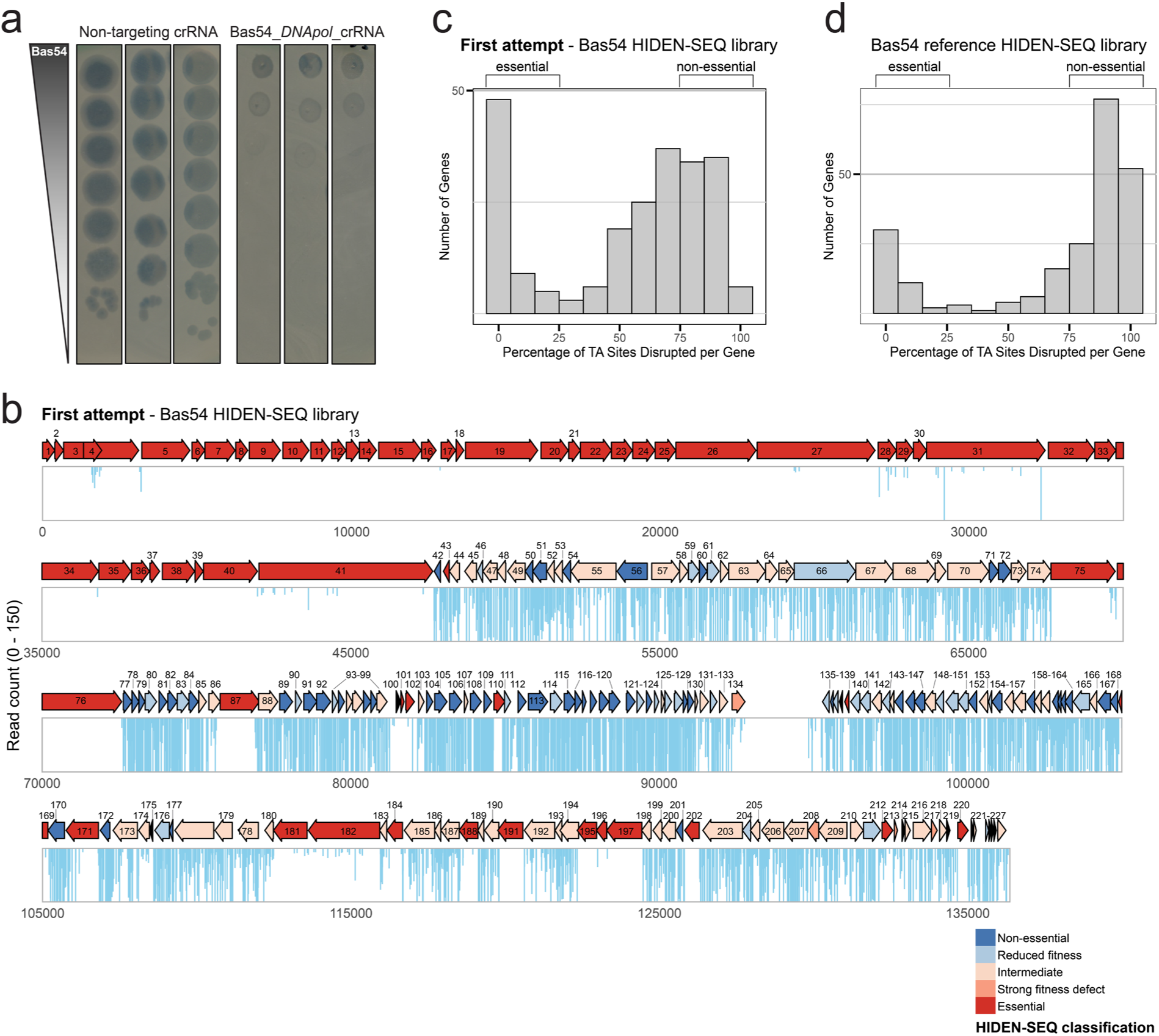
Genome-wide essentiality mapping of Bas54 phage. **a**, Ten-fold serial dilution plaque assays of phage Bas54 plated on *E. coli* K-12 ΔRM expressing *cas13a* with either a non-targeting crRNA control or a crRNA targeting the transcripts of the DNA polymerase gene. **b**, Phage Bas54 HIDEN-SEQ gene essentiality map from the first library which was not highly saturated. The start of the genome map has been set to the start of the i-spanin gene at the beginning of the terminase operon. Arrows present the coding sequences with the orientation indicating the direction of transcription. Arrow colours correspond to HIDEN-SEQ classification categories derived from transposon insertion profiles: non-essential (dark blue), reduced fitness (light blue), intermediate (light orange), strong fitness defect (orange), and essential (red). Vertical bars represent the number of transposon insertions at TA dinucleotide sites with heights indicating read counts (up to a maximum of 150). Genes containing ≤5 TA sites are shown in black and were not classified. **c**, The histogram illustrates the percentage of disrupted TA sites per gene for the Bas54 HIDEN-SEQ first library showing many genes with multiple insertions but overall low coverage of TA sites. **d**, The histogram illustrates the percentage of disrupted TA sites per gene for the Bas54 reference HIDEN-SEQ library showing that non-essential and essential genes are easily distinguishable in a highly saturated library.

**Extended Data Fig. 6.**
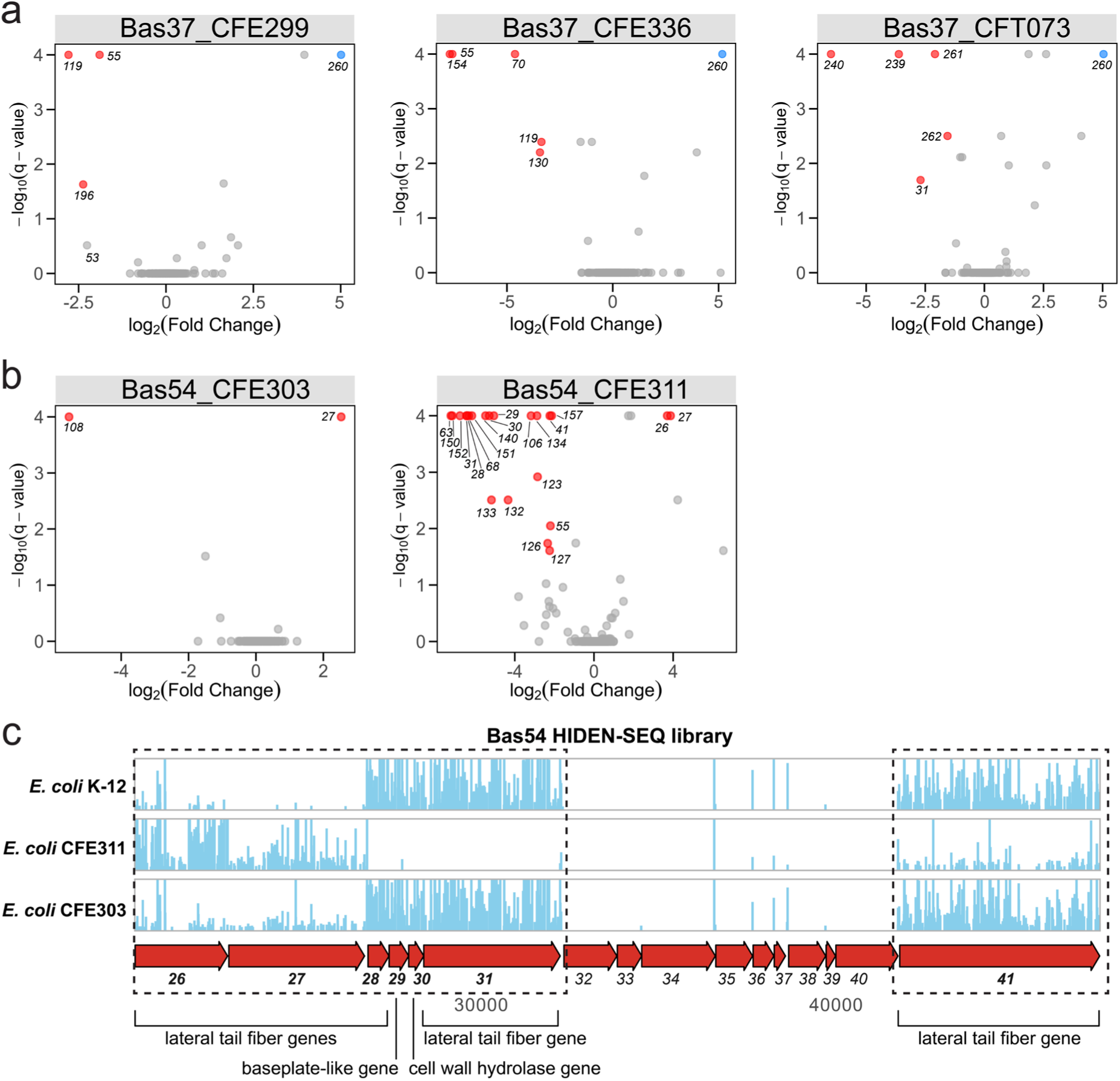
Identification of additional conditionally essential genes in clinical isolates by HIDEN-SEQ. **a**, Volcano plots comparing the depletion of gene disruptions in phage Bas37 after infection of the laboratory strain *E. coli* K-12 BW25113 with infections of uropathogenic isolates *E. coli* CFE299, CFE336, and CFT073. Each point represents a viral gene with the *x*-axis showing the log₂(fold change) in normalized insertion counts and the *y*-axis showing the –log₁₀(*q*-value) (see *Methods*). Negative log₂(fold change) values indicate that mutants with insertions in that gene had reduced fitness in the clinical isolate compared to the laboratory strain. Genes highlighted in red represent the most significant observed effects. *bas37_0119* encodes the Cef ortholog, *bas37_0196* is a gene of unknown function and NrdC.1 ortholog, *bas37_0055* encodes PseT ortholog. In all plots, the single blue point represents disruptions in *bas37_0260* which causes a strong fitness defect in *E. coli* K-12 BW25113 but not in the clinical isolates. **b**, Volcano plots comparing the depletion of gene disruptions in phage Bas54 after infection of the laboratory strain *E. coli* K-12 BW25113 with infections of uropathogenic isolates *E. coli* CFE303 or CFE311. Each point represents a viral gene with the *x*-axis showing the log₂(fold change) in normalized insertion counts and the *y*-axis *showing* the –log₁₀(*q*-value) (see *Methods*). Negative log₂(fold change) values indicate that mutants with insertions in that gene had reduced fitness in the clinical isolate compared to the laboratory strain. Genes highlighted in red represent the most significant observed effects. **c**, Transposon insertion profiles at the locus encoding lateral tail fiber genes of Bas54 obtained after infection of *E. coli* K-12 BW25113, *E. coli* CFE311, and *E. coli* CFE303 with the Bas54 reference HIDEN-SEQ library. Vertical bars represent the number of transposon insertions at TA sites with heights indicating read counts (up to a maximum of 150). It is apparent that different sets of lateral tail fiber genes are conditionally essential in different strains.

**Extended Data Fig. 7.**
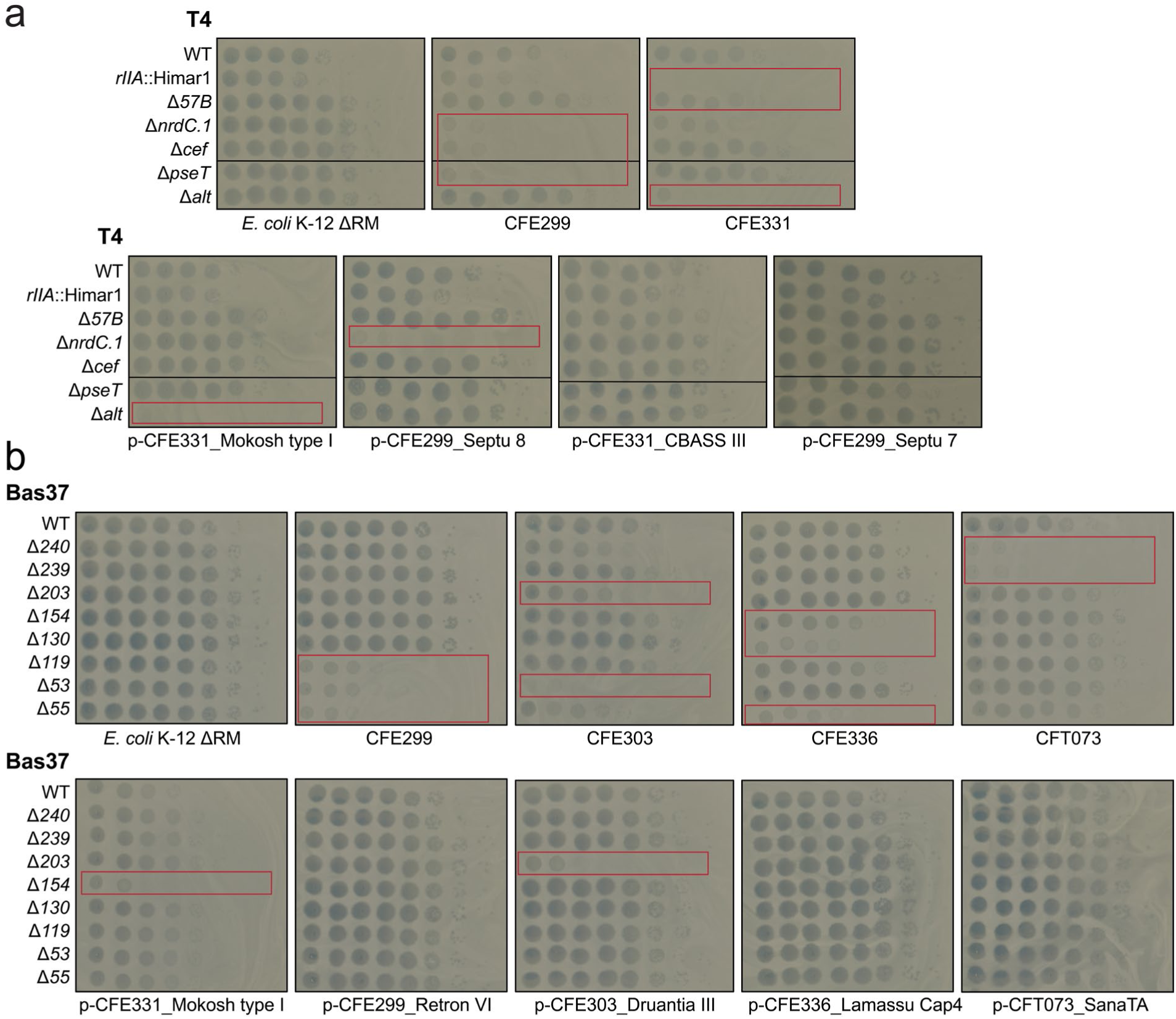
Matching candidate phage anti-defense genes identified by HIDEN-SEQ with host defense systems. The figure shows ten-fold serial dilution plaque assays of knockout mutants of phages T4 (**a**) and Bas37 (**b**) lacking candidate anti-defense genes plated on *E. coli* K-12 ΔRM variants expressing predicted defense systems from uropathogenic *E. coli* strains. In parallel, the same viral knockout mutants were also spotted on the uropathogenic strains to confirm the fitness defects detected by HIDEN-SEQ (see figS. 4a and 4c as well as Extended Data Figures 6a and 6b). Data are representative of 3 independent biological replicates.

**Extended Data Fig. 8.**
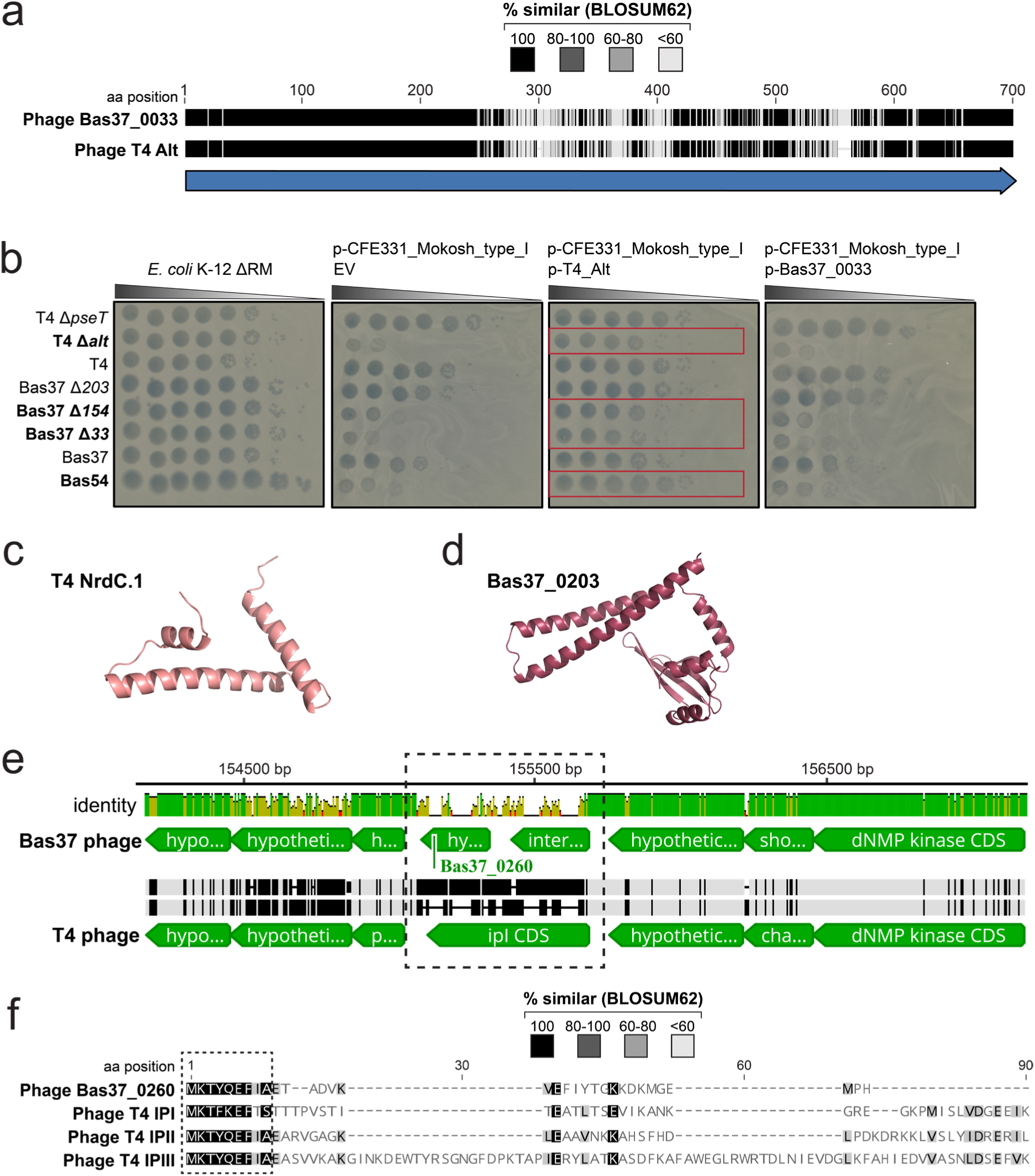
New anti-defense genes with known and unknown functions. **a**, Amino acid sequence alignment of Alt ADP-ribosyltransferase of phage Bas37 (encoded by *bas37_0033*) and phage T4 generated using MAFFT v7.490 implemented in Geneious Prime 2023.0.4 with default settings. **b**, Ten-fold serial dilution plaque assays of knockout mutants of phages T4 and Bas37 plated on *E. coli* K-12 ΔRM expressing the Mokosh type I with either an empty vector (EV) or expressing the Alt orthologs of T4 or Bas37 (encoded by *bas37_0033*). Plates show infections after induction with 5mM IPTG with T4 Δ*alt*, Bas37 Δ*33*, Bas37 Δ*154* (which lacks β*α*-glucosyltransferase), and the Mokosh-sensitive phage Bas54. Wildtype phages and knockout mutants T4 Δ*pseT* and Bas37 Δ*203* are shown as controls. Data are representative of 3 independent biological replicates. **c**, AlphaFold2-predicted structure of NrdC.1 of phage T4. **d**, AlphaFold2 predicted structure of Bas37_0203 of phage Bas37. **e**, The illustration shows a sequence alignment at the region comprising the *ipI* of phage T4. Note that the upstream and downstream regions are highly conserved. T4 contains the *ipI* gene, whereas Bas37 encodes two different genes in this locus, including *bas37_0260*. Alignments were generated using MAFFT v7.490 implemented in Geneious Prime 2023.0.4 with default settings. **f**, Partial amino acid sequence alignment of Bas37_0260 and phage T4 internal proteins IPI, IPII, IPIII, highlighting the conserved sequence motif at the N-terminal.

**Extended Data Table 1.** List of all bacterial strains used in this study.

**Extended Data Table 2.** List of all phages used in this study and annotation tables.

**Extended Data Table 3.** Selected defense systems predicted in *E. coli* uropathogenic isolates (using DefenseFinder).

**Extended Data Table 4.** Construction of all plasmids generated in this study.

**Extended Data Table 5.** List of all oligonucleotide primers and synthesized DNA used in this study.

**Extended Data Table 6.** Gene essentiality classification and conditional essentiality for phage T4.

**Extended Data Table 7.** Gene essentiality classification and conditional essentiality for phage Bas37.

**Extended Data Table 8.** Gene essentiality classification and conditional essentiality for phage Bas54.

