## Supplementary Information for "Systematic mapping of bacteriophage gene essentiality with HIDEN-SEQ"

### Supplementary Discussion

#### Phage anti-defense factors against Mokosh type I and regrowth-associated fitness effects

The ADP-ribosyltransferases and glucosyltransferases of T-even phages were identified as anti-defense factors against Mokosh type I system of *E. coli* CFE331 strain. Comparison between two closely related phages, T4 and Bas37, highlights overlapping mechanisms yet distinct gene requirements in their counter-defense against Mokosh (Fig. 4c). In Bas37, *alt*, *βα-gt*, and *modA* (*bas37_0131*), which is another known ADP-ribosyltransferase[71], were identified as conditionally essential genes. The requirements of Bas37 for both *modA* and *alt* suggest functional compensation for the weaker activity of Bas37 Alt (Extended Data Fig. 8b). In contrast, in T4 phage, complete loss of insertions in the presence of Mokosh type I was observed only for T4 *alt* and *α-gt* gene, while there was only a minor reduction in a few other genes, such as *modA*, *alc* and *β-gt*. These results also point to differences in glycosylation of hydroxymethyl cytosine residues, as previously reported for T-even relatives[72], which may also account for the higher sensitivity of Bas37 to Mokosh type I (Fig. 4b). Notably, the Bas37 *βα-gt* is orthologous to the T6 *βα-gt* gene, which adds a second glucosyl group to generate gentiobiosyl-hydroxymethyl cytosine[72]. However, the Bas37 *α-gt* gene did not appear conditionally essential in this case because its depletion was due to a regrowth-associated fitness effect rather than only selective pressure. Such effects were also observed for a few genes in other experiments where insertions tolerated in the reference library were progressively depleted upon passaging. Consequently, such genes may not be identified as conditionally essential. Similar effects have also been reported in bacterial TnSeq[73], where the measured fitness of insertions can shift across serial passages, indicating a gradual fitness disadvantage that is only revealed under longer selection. Such progressive changes can occur both broadly (Supplementary Fig. 1a, b) but also in specific phage-host settings (Supplementary Fig. 1c).

#### References (Supplementary Discussion)


71. Tiemann, B., Depping, R. & Rüger, W. Overexpression, Purification, and Partial Characterization of ADP-Ribosyltransferases ModA and ModB of Bacteriophage T4. *Gene Expression* **8**, 187 (2018).

72. Lehman, I. R. & Pratt, E. A. On the Structure of the Glucosylated Hydroxymethylcytosine Nucleotides of Coliphages T2, T4, and T6. *Journal of Biological Chemistry* **235**, 3254–3259 (1960).

73. Miravet-Verde, S., Burgos, R., Garcia-Ramallo, E., Weber, M. & Serrano, L. Quantitative essentiality in a reduced genome: a functional, regulatory and structural fitness map. *Molecular Systems Biology* 1–29 (2025) doi:10.1038/s44320-025-00133-1.

### Supplementary Figures

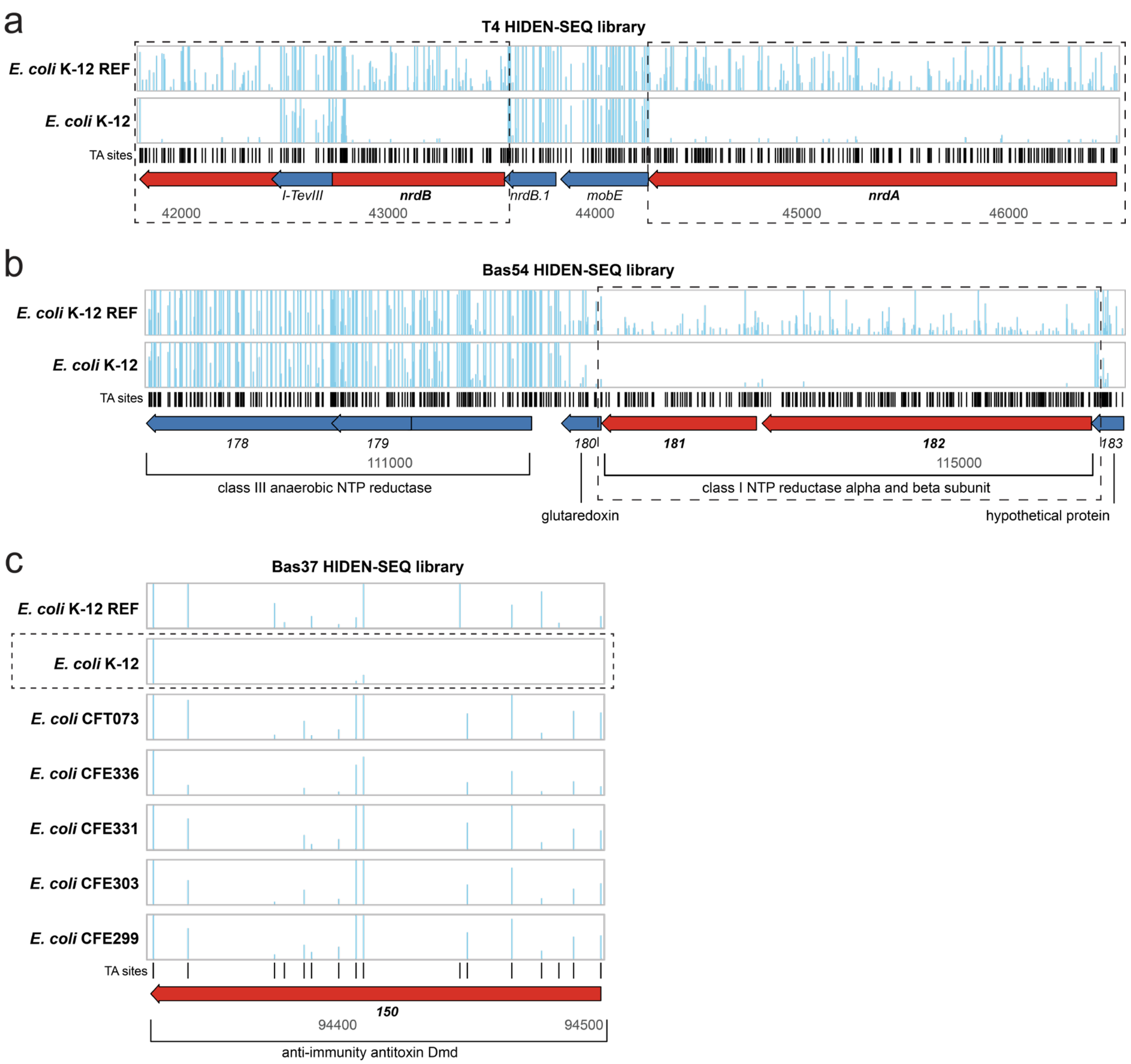


**Supplementary Fig. 1 | Gene fitness effects upon passaging of the transposon phage libraries.**

**a**, Transposon insertion profiles at the locus encoding ribonucleoside reductase subunits (*nrdA* and *nrdB)* genes of phage T4 for the T4 reference HIDEN-SEQ library (top track) and those obtained upon passaging of the library in *E. coli* K-12 BW25113 host (bottom track). Vertical bars represent the number of transposon insertions at TA sites (shown with black tick marks) with heights indicating read counts (up to a maximum of 150). The illustration highlights that the observed depletion of transposon insertion mutants is specific to *nrdA* and *nrdB* genes. This depletion was also observed in other hosts (Extended Data Table 6). **b**, Same as in (a), but for ribonucleotide reductase subunits in Bas54 phage. **c**, Transposon insertion profiles at the locus encoding the Dmd antitoxin (*bas37_0150*) gene ortholog of Bas37 phage of the Bas37 reference HIDEN-SEQ library (top track) and those obtained upon passaging of the library in the *E. coli* K-12 BW25113 host expressing *rnlAB* or different *E. coli* clinical isolates. The illustration highlights that the insertion depletion is host-specific because *bas37_0150* insertions are progressively depleted (more essential) upon passaging in *E. coli* K-12, but not in the other *E. coli* strains.
