## Extended Data Table 1 for "Systematic mapping of bacteriophage gene essentiality with HIDEN-SEQ"

### **Extended Data Table 1. List of all bacterial strains used in this study**

| **Identifier** | **Genotype** | **relevant plasmid(s)** | **Selection** | **Source/Description** |
| --- | --- | --- | --- | --- |
| AH-E03-200 | *E. coli* K-12 MG1655 ΔRM  MG1655 Δ*mrr-hsdRMS-mcrBC* Δ*mcrA* | none | none | Our laboratory collection^1^; strain lacking all known restriction systems of *E. coli* K-12 |
| AH-E01-047 | *E. coli* K-12 BW25113  F^-^ Δ(*araD-araB*)567, Δ*lacZ4787(::rrnB-3*), λ^-^, *rph-1*, Δ(*rhaD-rhaB*)568, *hsdR514* | none | none | Our laboratory collection^2^ |
| DH-E01-007 | *E. coli* K-12 MG1655 ΔRM  MG1655 Δ*mrr-hsdRMS-mcrBC* Δ*mcrA* | pAH221_LbuCas13a | Cam25 | Our laboratory collection; strain carrying the LbuCas13a expression plasmid |
| AH-E10-748 | *E. coli* K-12 DH5α *(?)* F*- endA1 glnV44 thi-1 recA1 relA1 gyrA96 deoR nupG Φ80dlacZΔM15 Δ(lacZYA-argF)U169, hsdR17(rK- mK+), λpir* | pAH218_LbuCas13a_e | Gen20 | Our laboratory collection; strain carrying crRNA cassette plasmid for cloning LbuCas13a spacers |
| DH-E01-008 | *E. coli* K-12 MG1655 ΔRM  MG1655 Δ*mrr-hsdRMS-mcrBC* Δ*mcrA* | pAH221_LbuCas13a, pAH218_LbuCas13a_e | Cam25, Gen20 | Our laboratory collection; strain carrying the LbuCas13a expression plasmid and the empty crRNA cassette plasmid |
| DH-E03-201 | *E. coli* K-12 MG1655 ΔRM  MG1655 Δ*mrr-hsdRMS-mcrBC* Δ*mcrA* | pAH221_LseCas13a | Cam25 | This study; strain carrying the LseCas13a expression plasmid |
| DH-E03-182 | *E. coli* K-12 DH5α *(?)* F*- endA1 glnV44 thi-1 recA1 relA1 gyrA96 deoR nupG Φ80dlacZΔM15 Δ(lacZYA-argF)U169, hsdR17(rK- mK+), λpir* | pAH218_LseCas13a_e | Gen20 | This study; strain carrying crRNA cassette plasmid for cloning LseCas13a spacers |
| DH-E03-173 | *E. coli* K-12 MG1655 ΔRM  MG1655 Δ*mrr-hsdRMS-mcrBC* Δ*mcrA* | pAH221_LseCas13a, pAH218_LseCas13a_e | Cam25, Gen20 | This study; strain carrying the LseCas13a expression plasmid and the empty crRNA cassette plasmid |
| DH-E02-160 | *E. coli* K-12 MG1655 ΔRM  MG1655 Δ*mrr-hsdRMS-mcrBC* Δ*mcrA* | pAH221_LbuCas13a, pAH218_LbuCas13a_Bas54_DNA_pol_H5 | Cam25, Gen20 | This study; LbuCas13a selection strain with crRNA targeting Bas54 phage |
| DH-E03-226 | *E. coli* K-12 MG1655 ΔRM  MG1655 Δ*mrr-hsdRMS-mcrBC* Δ*mcrA* | pAH221_LseCas13a,  pAH218_LseCas13a_2_T4_mcp_CDS_2 | Cam25, Gen20 | This study; LseCas13a selection strain with crRNA targeting T4/Bas37 phage |
| DH-E05-364 | *E. coli* K-12 BW25113  F^-^ Δ(*araD-araB*)567, Δ*lacZ4787(::rrnB-3*), λ^-^, *rph-1*, Δ(*rhaD-rhaB*)568, *hsdR514* | pAH186ColE1_Himar1_T'ase, pAH160_Tn_AIcrVIA3_v2 | Amp50, Kan50 | This study; transposition strain with transposon carrying AIcrVIA3 |
| DH-E04-282 | *E. coli* K-12 BW25113  F^-^ Δ(*araD-araB*)567, Δ*lacZ4787(::rrnB-3*), λ^-^, *rph-1*, Δ(*rhaD-rhaB*)568, *hsdR514* | pAH186ColE1_Himar1_T'ase, pDH160_Tn_AcrVIA1_v2 | Amp50, Kan50 | This study; transposition strain with transposon carrying AcrVIA1 |
| DH-E04-253 | *E. coli* K-12 BW25113  F^-^ Δ(*araD-araB*)567, Δ*lacZ4787(::rrnB-3*), λ^-^, *rph-1*, Δ(*rhaD-rhaB*)568, *hsdR514* | pAH186ColE1_Himar1_T'ase,  pAH160_Tn_sup-tRNAs_v2 | Amp50, Kan50 | This study; transposition strain with transposon carrying sup-tRNAs as non-selectable (with Cas13a) cassette |
| DH-E05-351 | *E. coli* K-12 BW25113  F^-^ Δ(*araD-araB*)567, Δ*lacZ4787(::rrnB-3*), λ^-^, *rph-1*, Δ(*rhaD-rhaB*)568, *hsdR514* | pSH232_EcCdnG-WT | Amp50 | Plasmid obtained from Prof. Philip J. Kranzusch; strain expressing CBASS EcCdnG operon^3^ |
| DH-E05-352 | *E. coli* K-12 BW25113  F^-^ Δ(*araD-araB*)567, Δ*lacZ4787(::rrnB-3*), λ^-^, *rph-1*, Δ(*rhaD-rhaB*)568, *hsdR514* | pSH233_EcCdnG-delHNH | Amp50 | Plasmid obtained from Prof. Philip J. Kranzusch; strain expressing CBASS EcCdnG operon with inactive Cap effector proteins^3^ |
| DH-E05-401 | *E. coli* K-12 BW25113  F^-^ Δ(*araD-araB*)567, Δ*lacZ4787(::rrnB-3*), λ^-^, *rph-1*, Δ(*rhaD-rhaB*)568, *hsdR514* | pBR322_darTG1 | Amp50 | Plasmid obtained from Prof. Michele LeRoux; strain expressing DarTG1 system^4^ |
| DH-E05-402 | *E. coli* K-12 BW25113  F^-^ Δ(*araD-araB*)567, Δ*lacZ4787(::rrnB-3*), λ^-^, *rph-1*, Δ(*rhaD-rhaB*)568, *hsdR514* | pBR322_darTG2 | Amp50 | Plasmid obtained from Prof. Michele LeRoux; strain expressing DarTG2 system^4^ |
| DH-E05-400 | *E. coli* K-12 BW25113  F^-^ Δ(*araD-araB*)567, Δ*lacZ4787(::rrnB-3*), λ^-^, *rph-1*, Δ(*rhaD-rhaB*)568, *hsdR514* | pBR322_ev | Amp50 | Plasmid obtained from Dr. Michele LeRoux; control strain for experiments with DarTG1 and DarTG2 carrying the empty vector^4^ |
| DH-E05-338 | *E. coli* K-12 BW25113  F^-^ Δ(*araD-araB*)567, Δ*lacZ4787(::rrnB-3*), λ^-^, *rph-1*, Δ(*rhaD-rhaB*)568, *hsdR514* | pUA139 | Kan50 | Our laboratory collection; control strain for experiments with RexAB carrying the empty vector |
| DH-E05-339 | *E. coli* K-12 BW25113  F^-^ Δ(*araD-araB*)567, Δ*lacZ4787(::rrnB-3*), λ^-^, *rph-1*, Δ(*rhaD-rhaB*)568, *hsdR514* | pVD3_RexAB_v2 | Kan50 | This study; strain expressing RexAB |
| DH-E04-321 | *E. coli* K-12 *ΔrnlA* BW25113 *rnlA::kanR(*pKD13*)* | none | Kan50 | Keio collection mutant^5^ obtained from Prof. Urs Jenal; strain not expressing RnlAB defense system |
| DH-E05-329 | *E. coli* K-12 *ΔrnlA* BW25113 *rnlA::kanR(*pKD13*)* | pAH221_LseCas13a, pAH218_LseCas13a_2_empty | Kan50, Cam25, Gen20 | Keio collection mutant not expressing RnlAB defense system carrying the LseCas13a expression plasmid and the empty crRNA cassette plasmid |
| DH-E05-330 | *E. coli* K-12 *ΔrnlA* BW25113 *rnlA::kanR(*pKD13*)* | pAH221_LseCas13a, pAH218_LseCas13a_2_T4_crRNA_mcp_CDS_2 | Kan50, Cam25, Gen20 | Keio collection mutant not expressing RnlAB defense system LseCas13a selection strain with crRNA targeting T4 phage |
| DH-E05-336 | *E. coli* K-12 *ΔrnlA* BW25113 *rnlA::FRT* | none | Kan50 | Keio collection mutant^5^ not expressing RnlAB defense system; Kanamycin cassette was FLPped out using pCP20 |
| DH-E05-346 | *E. coli* K-12 *ΔrnlA* BW25113 *rnlA::FRT* | pAH186ColE1_Himar1_T'ase, pDH160_Tn_AcrVIA1_v2 | Kan50, Amp50 | Keio collection mutant not expressing RnlAB defense system transposition strain with transposon carrying AcrVIA1 |
| DH-E06-408 | *E. coli* K-12 MG1655 ΔRM  MG1655 Δ*mrr-hsdRMS-mcrBC* Δ*mcrA* | pAH210_T4_tk.4_AcrVIA1 | Kan50 | This study; Strain carrying the homologous recombination plasmid for removing *tk.4*/*adfM* gene from T4 phage |
| DH-E05-396 | *E. coli* K-12 MG1655 ΔRM  MG1655 Δ*mrr-hsdRMS-mcrBC* Δ*mcrA* | pAH218_T4_vs_AcrVIA1 | Gen20 | This study; Strain carrying the homologous recombination plasmid for removing *vs* gene from T4 phage |
| DH-E06-412 | *E. coli* K-12 MG1655 ΔRM  MG1655 Δ*mrr-hsdRMS-mcrBC* Δ*mcrA* | pAH210_T4_alt_AcrVIA1 | Kan50 | This study; Strain carrying the homologous recombination plasmid for removing *alt* gene from T4 phage |
| DH-E06-413 | *E. coli* K-12 MG1655 ΔRM  MG1655 Δ*mrr-hsdRMS-mcrBC* Δ*mcrA* | pAH210_T4_cef_AcrVIA1 | Kan50 | This study; Strain carrying the homologous recombination plasmid for removing *cef* gene from T4 phage |
| DH-E06-414 | *E. coli* K-12 MG1655 ΔRM  MG1655 Δ*mrr-hsdRMS-mcrBC* Δ*mcrA* | pAH210_T4_gp57B_AcrVIA1 | Kan50 | This study; Strain carrying the homologous recombination plasmid for removing *57B* gene from T4 phage |
| DH-E06-415 | *E. coli* K-12 MG1655 ΔRM  MG1655 Δ*mrr-hsdRMS-mcrBC* Δ*mcrA* | pAH210_T4_nrdC.1_AcrVIA1 | Kan50 | This study; Strain carrying the homologous recombination plasmid for removing *nrdC.1* gene from T4 phage |
| DH-E06-416 | *E. coli* K-12 MG1655 ΔRM  MG1655 Δ*mrr-hsdRMS-mcrBC* Δ*mcrA* | pAH210_T4_rnlA_AcrVIA1_1 | Kan50 | This study; Strain carrying the homologous recombination plasmid for inserting *acrVIA1* gene in T4 *rnlA* gene – position 1 |
| DH-E06-417 | *E. coli* K-12 MG1655 ΔRM  MG1655 Δ*mrr-hsdRMS-mcrBC* Δ*mcrA* | pAH210_T4_rnlA_AcrVIA1_2 | Kan50 | This study; Strain carrying the homologous recombination plasmid for inserting *acrVIA1* gene in T4 *rnlA* gene – position 2 |
| DH-E06-433 | *E. coli* K-12 MG1655 ΔRM  MG1655 Δ*mrr-hsdRMS-mcrBC* Δ*mcrA* | pAH210_T4_pseT_AcrVIA1 | Kan50 | This study; Strain carrying the homologous recombination plasmid for removing *pseT* gene from T4 phage |
| DH-E06-418 | *E. coli* K-12 MG1655 ΔRM  MG1655 Δ*mrr-hsdRMS-mcrBC* Δ*mcrA* | pAH210_Bas37_0053_AcrVIA1 | Kan50 | This study; Strain carrying the homologous recombination plasmid for removing *bas37_0053* gene from Bas37 phage |
| DH-E06-419 | *E. coli* K-12 MG1655 ΔRM  MG1655 Δ*mrr-hsdRMS-mcrBC* Δ*mcrA* | pAH210_Bas37_0055_AcrVIA1 | Kan50 | This study; Strain carrying the homologous recombination plasmid for removing *bas37_0055* gene from Bas37 phage |
| DH-E06-420 | *E. coli* K-12 MG1655 ΔRM  MG1655 Δ*mrr-hsdRMS-mcrBC* Δ*mcrA* | pAH210_Bas37_0119_AcrVIA1 | Kan50 | This study; Strain carrying the homologous recombination plasmid for removing *bas37_0119* gene from Bas37 phage |
| DH-E06-421 | *E. coli* K-12 MG1655 ΔRM  MG1655 Δ*mrr-hsdRMS-mcrBC* Δ*mcrA* | pAH210_Bas37_0130_AcrVIA1 | Kan50 | This study; Strain carrying the homologous recombination plasmid for removing *bas37_0130* gene from Bas37 phage |
| DH-E06-422 | *E. coli* K-12 MG1655 ΔRM  MG1655 Δ*mrr-hsdRMS-mcrBC* Δ*mcrA* | pAH210_Bas37_0154_AcrVIA1 | Kan50 | This study; Strain carrying the homologous recombination plasmid for removing *bas37_0154* gene from Bas37 phage |
| DH-E06-423 | *E. coli* K-12 MG1655 ΔRM  MG1655 Δ*mrr-hsdRMS-mcrBC* Δ*mcrA* | pAH210_Bas37_0203_AcrVIA1 | Kan50 | This study; Strain carrying the homologous recombination plasmid for removing *bas37_0203* gene from Bas37 phage |
| DH-E06-424 | *E. coli* K-12 MG1655 ΔRM  MG1655 Δ*mrr-hsdRMS-mcrBC* Δ*mcrA* | pAH210_Bas37_0239_AcrVIA1 | Kan50 | This study; Strain carrying the homologous recombination plasmid for removing *bas37_0239* gene from Bas37 phage |
| DH-E06-425 | *E. coli* K-12 MG1655 ΔRM  MG1655 Δ*mrr-hsdRMS-mcrBC* Δ*mcrA* | pAH210_Bas37_0240_AcrVIA1 | Kan50 | This study; Strain carrying the homologous recombination plasmid for removing *bas37_0240* gene from Bas37 phage |
| DH-E06-460 | *E. coli* K-12 MG1655 ΔRM  MG1655 Δ*mrr-hsdRMS-mcrBC* Δ*mcrA* | pAH210_Bas37_0033_AcrVIA1 | Kan50 | This study; Strain carrying the homologous recombination plasmid for removing *bas37_0033* gene from Bas37 phage |
| DH-E06-475 | *E. coli* K-12 MG1655 ΔRM  MG1655 Δ*mrr-hsdRMS-mcrBC* Δ*mcrA* | pAH210_Bas37_0260_AcrVIA1 | Kan50 | This study; Strain carrying the homologous recombination plasmid for removing *bas37_0260* gene from Bas37 phage |
| DH-E06-426 | *E. coli* K-12 MG1655 ΔRM  MG1655 Δ*mrr-hsdRMS-mcrBC* Δ*mcrA* | pAH210_Bas54_0063_AIcrVIA3 | Kan50 | This study; Strain carrying the homologous recombination plasmid for removing *bas54_0063* gene from Bas54 phage |
| DH-E06-428 | *E. coli* K-12 MG1655 ΔRM  MG1655 Δ*mrr-hsdRMS-mcrBC* Δ*mcrA* | pAH210_Bas54_0140_AIcrVIA3 | Kan50 | This study; Strain carrying the homologous recombination plasmid for removing *bas54_0140* gene from Bas54 phage |
| DP-E06-433 | *E. coli* K-12 MG1655 ΔRM  MG1655 Δ*mrr-hsdRMS-mcrBC* Δ*mcrA* | pAH210_CFE299_Septu_7 | Kan50 | This study; the pAH210 plasmid encoding antiviral defense from CFE299 clinical isolate |
| DP-E06-441 | *E. coli* K-12 MG1655 ΔRM  MG1655 Δ*mrr-hsdRMS-mcrBC* Δ*mcrA* | pAH210_CFE299_Retron_VI | Kan50 | This study; the pAH210 plasmid encoding antiviral defense from CFE299 clinical isolate |
| DP-E06-443 | *E. coli* K-12 MG1655 ΔRM  MG1655 Δ*mrr-hsdRMS-mcrBC* Δ*mcrA* | pAH210_CFE299_Septu_8 | Kan50 | This study; the pAH210 plasmid encoding antiviral defense from CFE299 clinical isolate |
| DP-E06-435 | *E. coli* K-12 MG1655 ΔRM  MG1655 Δ*mrr-hsdRMS-mcrBC* Δ*mcrA* | pAH210_CFE303_Druantia_III_1 | Kan50 | This study; the pAH210 plasmid encoding antiviral defense from CFE303 clinical isolate |
| DP-E06-449 | *E. coli* K-12 MG1655 ΔRM  MG1655 Δ*mrr-hsdRMS-mcrBC* Δ*mcrA* | pAH210_CFE331_CBASS_III_3 | Kan50 | This study; the pAH210 plasmid encoding antiviral defense from CFE331 clinical isolate |
| DP-E06-450 | *E. coli* K-12 MG1655 ΔRM  MG1655 Δ*mrr-hsdRMS-mcrBC* Δ*mcrA* | pAH210_CFE331_Mokosh_Type_I_A_7 | Kan50 | This study; the pAH210 plasmid encoding antiviral defense from CFE331 clinical isolate |
| DP-E06-437 | *E. coli* K-12 MG1655 ΔRM  MG1655 Δ*mrr-hsdRMS-mcrBC* Δ*mcrA* | pAH210_CFE336_Lamassu-Cap4_nuclease_1 | Kan50 | This study; the pAH210 plasmid encoding antiviral defense from CFE336 clinical isolate |
| DP-E06-439 | *E. coli* K-12 MG1655 ΔRM  MG1655 Δ*mrr-hsdRMS-mcrBC* Δ*mcrA* | pAH210_CFT073_SanaTA_8 | Kan50 | This study; the pAH210 plasmid encoding antiviral defense from CFT073 clinical isolate |
| DH-E06-469 | *E. coli* K-12 MG1655 ΔRM  MG1655 Δ*mrr-hsdRMS-mcrBC* Δ*mcrA* | pAH210_CFE311_DS-28_1 | Kan50 | This study; the pAH210 plasmid encoding antiviral defense from CFE311 clinical isolate |
| DH-E06-472 | *E. coli* K-12 MG1655 ΔRM  MG1655 Δ*mrr-hsdRMS-mcrBC* Δ*mcrA* | pAH210_CFE311_Gao_Tmn_3 | Kan50 | This study; the pAH210 plasmid encoding antiviral defense from CFE311 clinical isolate |
| DH-E06-462 | *E. coli* K-12 MG1655 ΔRM  MG1655 Δ*mrr-hsdRMS-mcrBC* Δ*mcrA* | pNDM220_T4_alt | Amp30 | This study; strain expressing the *alt* gene of T4 from a plasmid |
| DH-E06-461 | *E. coli* K-12 MG1655 ΔRM  MG1655 Δ*mrr-hsdRMS-mcrBC* Δ*mcrA* | pNDM220_Bas37_alt (0033) | Amp30 | This study; strain expressing the *alt* gene of Bas37 from a plasmid |
| DH-E06-463 | *E. coli* K-12 MG1655 ΔRM  MG1655 Δ*mrr-hsdRMS-mcrBC* Δ*mcrA* | pAH210_CFE331_Mokosh_Type_I_A_7, pNDM220 | Kan50, Amp30 | This study; control strain carrying pNDM220 empty plasmid and plasmid expressing Mokosh type I defense system from CFE331 clinical isolate |
| DH-E06-464 | *E. coli* K-12 MG1655 ΔRM  MG1655 Δ*mrr-hsdRMS-mcrBC* Δ*mcrA* | pAH210_CFE331_Mokosh_Type_I_A_7, pNDM220_T4_alt | Kan50, Amp30 | This study; strain expressing the *alt* gene of T4 from a plasmid and Mokosh type I defense system from CFE331 clinical isolate |
| DH-E06-465 | *E. coli* K-12 MG1655 ΔRM  MG1655 Δ*mrr-hsdRMS-mcrBC* Δ*mcrA* | pAH210_CFE331_Mokosh_Type_I_A_7, pNDM220_Bas37_alt(0033) | Kan50, Amp30 | This study; strain expressing the *alt* gene of Bas37 from a plasmid and Mokosh type I defense system from CFE331 clinical isolate |
| DH-E06-476 | *E. coli* K-12 MG1655 ΔRM  MG1655 Δ*mrr-hsdRMS-mcrBC* Δ*mcrA* | pAH213_mcrA | Amp50 | Our laboratory collection |
| DH-E06-477 | *E. coli* K-12 MG1655 ΔRM  MG1655 Δ*mrr-hsdRMS-mcrBC* Δ*mcrA* | pAH213_mcrBC | Amp50 | Our laboratory collection |
| DH-E06-478 | *E. coli* K-12 MG1655 ΔRM  MG1655 Δ*mrr-hsdRMS-mcrBC* Δ*mcrA* | pAH213_mrr | Amp50 | Our laboratory collection |
| AH-E04-284 | *E. coli* CFT073 *rpoS(+)* | none | none | Our laboratory collection |
| CFE-299 | *E. coli* isolate CFE299 | none | none | Collection of Swiss NCCR AntiResist consortium (to be published elsewhere) |
| CFE-303 | *E. coli* isolate CFE303 | none | none | Collection of Swiss NCCR AntiResist consortium (to be published elsewhere) |
| CFE-311 | *E. coli* isolate CFE311 | none | none | Collection of Swiss NCCR AntiResist consortium (to be published elsewhere) |
| CFE-331 | *E. coli* isolate CFE331 | none | none | Collection of Swiss NCCR AntiResist consortium (to be published elsewhere) |
| CFE-336 | *E. coli* isolate CFE336 | none | none | Collection of Swiss NCCR AntiResist consortium (to be published elsewhere) |

### **References (Extended Data Table 1)**
