## Extended Data Table 5 for "Systematic mapping of bacteriophage gene essentiality with HIDEN-SEQ"

### Extended Data Table 5. List of all oligonucleotide primers and synthesized DNA used in this study

| ***Name*** | ***Purpose*** | ***Sequence (5'-3')*** |
| --- | --- | --- |
| prAH2173 | Used for PCR amplification; see Extended Data Table 4 | GATCTGATAGAGAAGGGTTTGC |
| prAH2179 | Used for PCR amplification; see Extended Data Table 4 | GTCATTTCGAACCCCAGAGT |
| prDH0264 | Used for PCR amplification; see Extended Data Table 4 | GCGGGACTCTGGGGTTCGAAATGAC |
| prDH0265 | Used for PCR amplification; see Extended Data Table 4 | ACGAGCAAACCCTTCTCTATCAGATC |
| prDH0347 | Used for PCR amplification; see Extended Data Table 4 | TGGCTGATAAGTCCCCGGTCTACGAAGCATGAGACCGATAGGTAAACGA |
| prDH0349 | Used for PCR amplification; see Extended Data Table 4 | CTATCGGTCTCATGCTTAGACCGGGGACTTATCAGCCAA |
| prDH0263 | Used for PCR amplification; see Extended Data Table 4 | CTGACGACCGCGCAAGTGGCACTTT |
| prDH0454 | Used for PCR amplification; see Extended Data Table 4 | TGAGACCGATAGGTAAACGA |
| prDH0263 | Used for PCR amplification; see Extended Data Table 4 | CTGACGACCGCGCAAGTGGCACTTT |
| prDH0233 | Used for PCR amplification; see Extended Data Table 4 | TCCAGTCGGTAGATATTCCACAAAACAGC |
| prDH0261 | Used for PCR amplification; see Extended Data Table 4 | GCTGTTTTGTGGAATATCTACCGACTGG |
| prDH0262 | Used for PCR amplification; see Extended Data Table 4 | CCCGAAAAGTGCCACTTGCG |
| prDH0343 | Used for PCR amplification; see Extended Data Table 4 | TACCTAGGACTGAGCTAGCCGTAAATCTCGAGTCATTTCGAACCC |
| prDH0344 | Used for PCR amplification; see Extended Data Table 4 | TTTACGGCTAGCTCAGTCCTAGGTACAATGCTAGCGGATCCTCTAGATTTTTTAAGAAGGAGATATA |
| prAH2568 | Used for PCR amplification; see Extended Data Table 4 | CAACTTAATCGCCTTGCAGC |
| prAH2245 | Used for PCR amplification; see Extended Data Table 4 | TCCAATTGTTATCCGCTCAC |
| prAH2525 | Used for PCR amplification; see Extended Data Table 4 | CAATTGTGAGCGGATAACAATTGGACTATGACCATGATTACGAATTCG |
| prAH2569 | Used for PCR amplification; see Extended Data Table 4 | GATGTGCTGCAAGGCGATTAAGTTGGTATACACTCCGCTATCGCTACC |
| prJR19 | Used for PCR amplification; see Extended Data Table 4 | TAATTTGACCAGAGAACAAGAATAACACTTTTCGGGGAAATGTTCTA |
| prJR21 | Used for PCR amplification; see Extended Data Table 4 | GCATAGTCCCTAGGACTGAGCTAGCTGTCAAGTTAATGTCATGATAATAATGGTTTCTTAGAC |
| prJR22 | Used for PCR amplification; see Extended Data Table 4 | AGCTAGCTCAGTCCTAGGGACTATGCTAGCTAAGAAGGAGATATACATATGAAGAATGGTTTTTATGCG |
| prJR18 | Used for PCR amplification; see Extended Data Table 4 | TTATTCTTGTTCTCTGGTCAAATTA |
| prAH2038 | Used for PCR amplification; see Extended Data Table 4 | GAGAGAAGATTTTCAGCCTGATAC |
| prAH2711 | Used for PCR amplification; see Extended Data Table 4 | TGTATCAGGCTGAAAATCTTCTCTCCTGCAAGGTAGTGGACAAGAC |
| prDH0210 | Used for PCR amplification; see Extended Data Table 4 | CGCTAGTTTGTTATCAGAATCGCAGAGCGAGCTCGATATCAAATTAC |
| prDH0204 | Used for PCR amplification; see Extended Data Table 4 | AAGTAAAGTGATTAACAGCGCATTAGAGCTGCTTAATGAGGTC |
| prDH0205 | Used for PCR amplification; see Extended Data Table 4 | CTAATGCGCTGTTAATCACTTTAC |
| prDH0206 | Used for PCR amplification; see Extended Data Table 4 | CTATTGGAATTCACGGTACTTTCAATTTCAT |
| prDH0207 | Used for PCR amplification; see Extended Data Table 4 | AAATTGAAAGTACCGTGAATTCCAATAGC |
| prDH0208 | Used for PCR amplification; see Extended Data Table 4 | GACTGTAACAGCTCTTCAGATACATACAGAG |
| prDH0209 | Used for PCR amplification; see Extended Data Table 4 | GTATGTATCTGAAGAGCTGTTACAGTCCTTG |
| prDH0153 | Used for PCR amplification; see Extended Data Table 4 | CTGCGATTCTGATAACAAACTAGC |
| prAH2697 | Used for PCR amplification; see Extended Data Table 4 | GCGGGACTCTGGGGTTCGAAATGACGGAATTCTTGACAGCTAGCTCA |
| prAH2698 | Used for PCR amplification; see Extended Data Table 4 | CGAGCAAACCCTTCTCTATCAGATCCCTTTCGTTTTATTTGATGCC |
| prDH0292 | Used for cloning **crRNA** against T4/Bas37; see Extended Data Table 4 | AAAC**TTCAAAAGTTCAGCTTTAGTTTTG**A |
| prDH0293 | Used for cloning **crRNA** against T4/Bas37; see Extended Data Table 4 | AGCAT**CAAAACTAAAGCTGAACTTTTGAA** |
| prDH0184 | Used for cloning **crRNA** against Bas54; see Extended Data Table 4 | AAAC**AACTTCAAAATCCTTATAAAGAGTTTTCTTT**A |
| prDH0185 | Used for cloning **crRNA** against Bas54; see Extended Data Table 4 | AGCAT**AAAGAAAACTCTTTATAAGGATTTTGAAGTT** |
| prAH2224 | Used for PCR amplification; see Extended Data Table 4 | CCCTTTGATATGTAACGGTGAAC |
| prAH2225 | Used for PCR amplification; see Extended Data Table 4 | GTTAATGTCATGATAATAATGGTTTCTTAGAC |
| prDH0509 | Used for PCR amplification; see Extended Data Table 4 | GCAAAACCCGTACCCTAGGT |
| prDH0154 | Used for PCR amplification; see Extended Data Table 4 | TCGAGATTGACAGCTAGCTCA |
| prDH0523 | Used for PCR amplification; see Extended Data Table 4 | AAACCATTATTATCATGACATTAACCTTCGTGGATAAGAGAAAAGAGTG |
| prDH0521 | Used for PCR amplification; see Extended Data Table 4 | CCTAGACCTAGGGTACGGGTTTTGCGAAATTGTGATTGTTGATTGGG |
| prDH0522 | Used for PCR amplification; see Extended Data Table 4 | GGACTGAGCTAGCTGTCAATCTCGAGCCCTTGATATATTTTACAATCATCTA |
| prDH0524 | Used for PCR amplification; see Extended Data Table 4 | CTGTTCACCGTTACATATCAAAGGGTGCGTCAGCAGATACATCATATAC |
| prDH0507 | Used for PCR amplification; see Extended Data Table 4 | CGGGACTCTGGGGTTCGAAATGACTCACCCCATTCAGTCTGTAAT |
| prDH0508 | Used for PCR amplification; see Extended Data Table 4 | CCTAGACCTAGGGTACGGGTTTTGCTGGTAGATTCTTATAATCGAGGATTG |
| prDH0510 | Used for PCR amplification; see Extended Data Table 4 | GGACTGAGCTAGCTGTCAATCTCGACCTATACATAAAACCAAAATTTTAGTCAT |
| prDH0511 | Used for PCR amplification; see Extended Data Table 4 | CGAGCAAACCCTTCTCTATCAGATCCCTCTTATAATTCGGGATGGA |
| prDH0526 | Used for PCR amplification; see Extended Data Table 4 | AAACCATTATTATCATGACATTAACCACCTGTCATTGTAACCAATTCA |
| prDH0527 | Used for PCR amplification; see Extended Data Table 4 | CCTAGACCTAGGGTACGGGTTTTGCCATTTTGCCACGTGGATTAA |
| prDH0528 | Used for PCR amplification; see Extended Data Table 4 | GGACTGAGCTAGCTGTCAATCTCGAGCGGTATTTTCTTCTTTGGATATAAG |
| prDH0529 | Used for PCR amplification; see Extended Data Table 4 | CTGTTCACCGTTACATATCAAAGGGCATGATTGAATTTATAGATGCATATGC |
| prDH0530 | Used for PCR amplification; see Extended Data Table 4 | AAACCATTATTATCATGACATTAACGGCTCCATACCTAAACGTCG |
| prDH0531 | Used for PCR amplification; see Extended Data Table 4 | CCTAGACCTAGGGTACGGGTTTTGCATGAGAAAGTCGGTGATAAACAG |
| prDH0532 | Used for PCR amplification; see Extended Data Table 4 | GGACTGAGCTAGCTGTCAATCTCGATTTACGTTTCATTACAATTTCCTCA |
| prDH0533 | Used for PCR amplification; see Extended Data Table 4 | CTGTTCACCGTTACATATCAAAGGGTATAGCAATGCTCTCTATGCATAAAG |
| prDH0534 | Used for PCR amplification; see Extended Data Table 4 | AAACCATTATTATCATGACATTAACCCTCATCAGTATGTAAACAACTTTGT |
| prDH0535 | Used for PCR amplification; see Extended Data Table 4 | CCTAGACCTAGGGTACGGGTTTTGCTGTCTTATAATGTTGGGCCC |
| prDH0536 | Used for PCR amplification; see Extended Data Table 4 | GGACTGAGCTAGCTGTCAATCTCGAAAATTCCATCATTCTTCATCTTTTG |
| prDH0537 | Used for PCR amplification; see Extended Data Table 4 | CTGTTCACCGTTACATATCAAAGGGGATCAAGCTGCTCGTCTGAT |
| prDH0538 | Used for PCR amplification; see Extended Data Table 4 | AAACCATTATTATCATGACATTAACTAGTCAGAAGCTCAGCAATTTTCT |
| prDH0539 | Used for PCR amplification; see Extended Data Table 4 | CCTAGACCTAGGGTACGGGTTTTGCGACGAGACTTTTAAACAAGTACTAGGA |
| prDH0540 | Used for PCR amplification; see Extended Data Table 4 | GGACTGAGCTAGCTGTCAATCTCGACAGTCTCCATATATTCTTTACGCTTT |
| prDH0541 | Used for PCR amplification; see Extended Data Table 4 | CTGTTCACCGTTACATATCAAAGGGTGATGATTATAGCCCTCCTGTTA |
| prDH0546 | Used for PCR amplification; see Extended Data Table 4 | AAACCATTATTATCATGACATTAACTGCTTCCATTGCATAAGTCTTT |
| prDH0547 | Used for PCR amplification; see Extended Data Table 4 | CCTAGACCTAGGGTACGGGTTTTGCTGATGGTGCATCAGATGATC |
| prDH0548 | Used for PCR amplification; see Extended Data Table 4 | GGACTGAGCTAGCTGTCAATCTCGATAATAGTCTTAAACAATTTTTCTGGATTATC |
| prDH0549 | Used for PCR amplification; see Extended Data Table 4 | CTGTTCACCGTTACATATCAAAGGGTCTAGTTGAACGATACGAAATCG |
| prDH0550 | Used for PCR amplification; see Extended Data Table 4 | AAACCATTATTATCATGACATTAACAAGTCTATCACGCAACCGAT |
| prDH0551 | Used for PCR amplification; see Extended Data Table 4 | CCTAGACCTAGGGTACGGGTTTTGCTCGATTTAAACGATGTTGATTATAT |
| prDH0552 | Used for PCR amplification; see Extended Data Table 4 | GGACTGAGCTAGCTGTCAATCTCGATATTCATCGTGAACGGATTTTC |
| prDH0553 | Used for PCR amplification; see Extended Data Table 4 | CTGTTCACCGTTACATATCAAAGGGCTCAGATGATGTAAGTGCATCTG |
| prDH0542 | Used for PCR amplification; see Extended Data Table 4 | AAACCATTATTATCATGACATTAACCCAACTAAACCGCTATTTACAATAAC |
| prDH0543 | Used for PCR amplification; see Extended Data Table 4 | CCTAGACCTAGGGTACGGGTTTTGCGACGATGTAGTTAAAGAAGAAATTTTC |
| prDH0544 | Used for PCR amplification; see Extended Data Table 4 | GGACTGAGCTAGCTGTCAATCTCGACAGAACCAGGACAGCCAATA |
| prDH0545 | Used for PCR amplification; see Extended Data Table 4 | CTGTTCACCGTTACATATCAAAGGGCCAGATTCAGCGGTTAGAATC |
| prDH0554 | Used for PCR amplification; see Extended Data Table 4 | AAACCATTATTATCATGACATTAACGCTGCGTTATCAAGCTTTTC |
| prDH0555 | Used for PCR amplification; see Extended Data Table 4 | CCTAGACCTAGGGTACGGGTTTTGCCAGAAGACAAAGTAGATGGTTCAAC |
| prDH0556 | Used for PCR amplification; see Extended Data Table 4 | GGACTGAGCTAGCTGTCAATCTCGAAACAAAATAGATATTTCTTTTCTTTTTCAA |
| prDH0557 | Used for PCR amplification; see Extended Data Table 4 | CTGTTCACCGTTACATATCAAAGGGTTGTGTATTCACTAGTGAAATCAGAG |
| prDH0558 | Used for PCR amplification; see Extended Data Table 4 | AAACCATTATTATCATGACATTAACGGTCATCAATAGCTAATTTCACATC |
| prDH0559 | Used for PCR amplification; see Extended Data Table 4 | CCTAGACCTAGGGTACGGGTTTTGCAACCAAAGAAGACCCAACAA |
| prDH0560 | Used for PCR amplification; see Extended Data Table 4 | GGACTGAGCTAGCTGTCAATCTCGATAAAACCCAGGATTTTTAGCAATA |
| prDH0561 | Used for PCR amplification; see Extended Data Table 4 | CTGTTCACCGTTACATATCAAAGGGATACTCGCGAACTCAATGAAC |
| prDH0562 | Used for PCR amplification; see Extended Data Table 4 | AAACCATTATTATCATGACATTAACAAGGTTCCATACCTAAACGTCG |
| prDH0563 | Used for PCR amplification; see Extended Data Table 4 | CCTAGACCTAGGGTACGGGTTTTGCTATTTCGTGAAATTACTGAAGATGG |
| prDH0564 | Used for PCR amplification; see Extended Data Table 4 | GGACTGAGCTAGCTGTCAATCTCGACGTCATTAGTGCAGTTCTGAAC |
| prDH0565 | Used for PCR amplification; see Extended Data Table 4 | CTGTTCACCGTTACATATCAAAGGGTCGCAAAAATTGGGATTGTG |
| prDH0566 | Used for PCR amplification; see Extended Data Table 4 | AAACCATTATTATCATGACATTAACCCGTCTTTGTCTCGGTAAAC |
| prDH0567 | Used for PCR amplification; see Extended Data Table 4 | CCTAGACCTAGGGTACGGGTTTTGCAGTTGTCAGAAGCAGAAATGC |
| prDH0568 | Used for PCR amplification; see Extended Data Table 4 | GGACTGAGCTAGCTGTCAATCTCGATCTTTAGAACGTCTTTTTGCAGAT |
| prDH0569 | Used for PCR amplification; see Extended Data Table 4 | CTGTTCACCGTTACATATCAAAGGGGCATTTTTCATAAAGTTGTTTACAAG |
| prDH0570 | Used for PCR amplification; see Extended Data Table 4 | AAACCATTATTATCATGACATTAACAGCTTTAATCAAACGGGATTTT |
| prDH0571 | Used for PCR amplification; see Extended Data Table 4 | CCTAGACCTAGGGTACGGGTTTTGCTGAAGAAGACGAGCCATTAAAG |
| prDH0572 | Used for PCR amplification; see Extended Data Table 4 | GGACTGAGCTAGCTGTCAATCTCGAAACATATCAAGGGCAGAAACTG |
| prDH0573 | Used for PCR amplification; see Extended Data Table 4 | CTGTTCACCGTTACATATCAAAGGGTCGTCAGTATAAAGCAGGTTCTATT |
| prDH0574 | Used for PCR amplification; see Extended Data Table 4 | AAACCATTATTATCATGACATTAACCATTAATTGCGTAATAACCATATGC |
| prDH0575 | Used for PCR amplification; see Extended Data Table 4 | CCTAGACCTAGGGTACGGGTTTTGCGGAAGTTTGAAGAACGGTTAAAG |
| prDH0576 | Used for PCR amplification; see Extended Data Table 4 | GGACTGAGCTAGCTGTCAATCTCGAAAATTTGTCCTCCATCAACTGT |
| prDH0577 | Used for PCR amplification; see Extended Data Table 4 | CTGTTCACCGTTACATATCAAAGGGTACAAGACTGTTCCTCTGTGGTATT |
| prDH0578 | Used for PCR amplification; see Extended Data Table 4 | AAACCATTATTATCATGACATTAACCACATCAGTAGAAATTATCTGACGC |
| prDH0579 | Used for PCR amplification; see Extended Data Table 4 | CCTAGACCTAGGGTACGGGTTTTGCCGAAGCATGAAAATAAACGTATTC |
| prDH0580 | Used for PCR amplification; see Extended Data Table 4 | GGACTGAGCTAGCTGTCAATCTCGATTTACCTCTAGAAAATCTTCCGG |
| prDH0581 | Used for PCR amplification; see Extended Data Table 4 | CTGTTCACCGTTACATATCAAAGGGGGAGTAAATAGGAATGAAAATGAAA |
| prDH0582 | Used for PCR amplification; see Extended Data Table 4 | AAACCATTATTATCATGACATTAACTCAAACATTGAATTGAAATCACTTT |
| prDH0583 | Used for PCR amplification; see Extended Data Table 4 | CCTAGACCTAGGGTACGGGTTTTGCCTTGCTCGAGTTGAAACCTT |
| prDH0584 | Used for PCR amplification; see Extended Data Table 4 | GGACTGAGCTAGCTGTCAATCTCGACAGCAGTATACTGTTCTTCGACAG |
| prDH0585 | Used for PCR amplification; see Extended Data Table 4 | CTGTTCACCGTTACATATCAAAGGGCAAAAGGCGATGTTGAAATA |
| prDH0634 | Used for PCR amplification; see Extended Data Table 4 | AAACCATTATTATCATGACATTAACATTTATCCTTGAACGAACTTGTAAG |
| prDH0635 | Used for PCR amplification; see Extended Data Table 4 | CCTAGACCTAGGGTACGGGTTTTGCAGAGTTGACTTTGCATCATTTG |
| prDH0636 | Used for PCR amplification; see Extended Data Table 4 | GGACTGAGCTAGCTGTCAATCTCGAAGGTGCATAAACGAAAGCCT |
| prDH0637 | Used for PCR amplification; see Extended Data Table 4 | CTGTTCACCGTTACATATCAAAGGGCTAGATTATTAAAGGCCTTCGGG |
| prDH0646 | Used for PCR amplification; see Extended Data Table 4 | AAACCATTATTATCATGACATTAACTTTTCTGCTGTAGTCAATAATCCAC |
| prDH0647 | Used for PCR amplification; see Extended Data Table 4 | CCTAGACCTAGGGTACGGGTTTTGCGGGCAGCAGAAAATAAATCAA |
| prDH0648 | Used for PCR amplification; see Extended Data Table 4 | GGACTGAGCTAGCTGTCAATCTCGATTTCATTTTGATTTCCATTTGGT |
| prDH0649 | Used for PCR amplification; see Extended Data Table 4 | CTGTTCACCGTTACATATCAAAGGGTGAAGCTGAGCGTCTTTTCT |
| prDH0586 | Used for PCR amplification; see Extended Data Table 4 | AAACCATTATTATCATGACATTAACCAACGTCATAGCCGATCTGT |
| prDH0587 | Used for PCR amplification; see Extended Data Table 4 | GTCATTTCGAACCCCAGAGTCCCGCCTTCTACAGGGATAGGTAGCCC |
| prDH0588 | Used for PCR amplification; see Extended Data Table 4 | ATCTGATAGAGAAGGGTTTGCTCGTAACCGAGATTTTGTATTCTCAGTCT |
| prDH0589 | Used for PCR amplification; see Extended Data Table 4 | CTGTTCACCGTTACATATCAAAGGGCCTTAAGCTTTAGCCAAAATGG |
| prDH0598 | Used for PCR amplification; see Extended Data Table 4 | AAACCATTATTATCATGACATTAACGCTACGCTCGAAAGAGAGACT |
| prDH0599 | Used for PCR amplification; see Extended Data Table 4 | ATCTGATAGAGAAGGGTTTGCTCGTTTCAGCTACTCCAACCTTGATAA |
| prDH0600 | Used for PCR amplification; see Extended Data Table 4 | GTCATTTCGAACCCCAGAGTCCCGCGGAGTATTTAGCCGCCTTTG |
| prDH0601 | Used for PCR amplification; see Extended Data Table 4 | CTGTTCACCGTTACATATCAAAGGGTTCTTGCCAAACTTGACACTG |
| prDP0266 | Used for PCR amplification; see Extended Data Table 4 | AAACCATTATTATCATGACATTAACGCAAAACAATATCGCGTCC |
| prDP0267 | Used for PCR amplification; see Extended Data Table 4 | CTGTTCACCGTTACATATCAAAGGGTGGATGTTCTCAGCCTGTC |
| prDP0280 | Used for PCR amplification; see Extended Data Table 4 | AAACCATTATTATCATGACATTAACACTTCTTAGCGGTGATTCTG |
| prDP0281 | Used for PCR amplification; see Extended Data Table 4 | CTGTTCACCGTTACATATCAAAGGGGAGACCTTCGAGTTATCAATC |
| prDP0284 | Used for PCR amplification; see Extended Data Table 4 | AAACCATTATTATCATGACATTAACTGGTAACGCGATCATCAAG |
| prDP0285 | Used for PCR amplification; see Extended Data Table 4 | CTGTTCACCGTTACATATCAAAGGGTAAAGGATGGAGTAAAGTAGCG |
| prDP0270 | Used for PCR amplification; see Extended Data Table 4 | AAACCATTATTATCATGACATTAACGAGGATGTGGAGTGAGGGG |
| prDP0271 | Used for PCR amplification; see Extended Data Table 4 | CTGTTCACCGTTACATATCAAAGGGTTTATAGAAAATACCGCTCCCG |
| prDP0296 | Used for PCR amplification; see Extended Data Table 4 | AAACCATTATTATCATGACATTAACATTGAGTCAGGCAGCATAAG |
| prDP0297 | Used for PCR amplification; see Extended Data Table 4 | CTGTTCACCGTTACATATCAAAGGGCAAATGTACCGGAATTGGAG |
| prDP0298 | Used for PCR amplification; see Extended Data Table 4 | AAACCATTATTATCATGACATTAACGAACGATTACATCAGTATCATACC |
| prDP0299 | Used for PCR amplification; see Extended Data Table 4 | CTGTTCACCGTTACATATCAAAGGGCTCATTTGACATGGTTAACG |
| prDP0274 | Used for PCR amplification; see Extended Data Table 4 | AAACCATTATTATCATGACATTAACAAATACGCCCACGAAGTCCC |
| prDP0275 | Used for PCR amplification; see Extended Data Table 4 | CTGTTCACCGTTACATATCAAAGGGAAAAATGTTAGGCCTGCAAGC |
| prDP0272 | Used for PCR amplification; see Extended Data Table 4 | AAACCATTATTATCATGACATTAACATTAGCAGACAGGTAATGTGG |
| prDP0273 | Used for PCR amplification; see Extended Data Table 4 | CTGTTCACCGTTACATATCAAAGGGCGGAATGAAACTGATTGTATAAC |
| prJR46 | Used for PCR amplification; see Extended Data Table 4 | AAACCATTATTATCATGACATTAACCATAATCCGGGTCAATGTAA |
| prJR47 | Used for PCR amplification; see Extended Data Table 4 | CTGTTCACCGTTACATATCAAAGGGGCAAGGCGGCTTTTATATAT |
| prJR48 | Used for PCR amplification; see Extended Data Table 4 | AAACCATTATTATCATGACATTAACGGCAATACTGGTGACGAATAA |
| prJR49 | Used for PCR amplification; see Extended Data Table 4 | CTGTTCACCGTTACATATCAAAGGGATTCTCTCCCGGGTAAGTGT |
| prDH0640 | Used for PCR amplification; see Extended Data Table 4 | ACAACGTCGTGACTGGGAAAAC |
| prDH0633 | Used for PCR amplification; see Extended Data Table 4 | AAAACGACGGCCAGTGAATTC |
| prDH0632 | Used for PCR amplification; see Extended Data Table 4 | CGAGGAATTCACTGGCCGTCGTTTTGCAGGAGGAATTCACCATGATGGAACTTATTACAGAATTATTTGAC |
| prDH0639 | Used for PCR amplification; see Extended Data Table 4 | AGGGTTTTCCCAGTCACGACGTTGTTTATCCTTGAACGAACTTGTAAGG |
| prAH1948 | Used for PCR amplification; see Extended Data Table 4 | GGGAACGCGGCCGCACCTACATCTGTATTAACGAAGCG, with NotI restriction site |
| prAH1949 | Used for PCR amplification; see Extended Data Table 4 | CTGCTTCTCGAGACACGGTGCCTGACTGC; with XhoI restriction site in the primer and adjacent ClaI + HindIII restriction sites on the amplified backbone |
| prAH1954 | Used for PCR amplification; see Extended Data Table 4 | CGAAACAAGCTTTTAAGGCCCATAACATCTCA; with HindIII restriction site |
| prAH1955 | Used for PCR amplification; see Extended Data Table 4 | GCATTTGCGGCCGCTCACTCAAAATAGTCCATATCCAG; NotI restriction site |
| prAH1958 | Used for PCR amplification; see Extended Data Table 4 | GGGAATATCGATGTAAACAGAAAGTCGCGTAACAG; with ClaI restriction site |
| prAH1959 | Used for PCR amplification; see Extended Data Table 4 | GCATTTGCGGCCGCTTATTTGAGATATTCATCGAAAATGT; with NotI restriction site |
| prAH1960 | Used for PCR amplification; see Extended Data Table 4 | GGGAACATCGATACAAAAGCTGGACTGGAAAC; with ClaI restriction site |
| prAH1961 | Used for PCR amplification; see Extended Data Table 4 | GCATTTGCGGCCGCTTATTTCTGTAATCGGTTTATATTAACG; with NotI restriction site |
| prDH0512 | To screen for T4 *vs* and *tk.4* knockout mutant | GAATAAACGATCCAATTCAGCTT |
| prDH0513 | To screen for T4 *vs* knockout mutant | CTATGCCTGCACCAATCCTA |
| prDH0318 | To screen for recombinant phages; anneals in *acrVIA1* | ACTTCTGTCTCGTTGTGCGT |
| prDH0614 | To screen for T4 *alt* knockout mutant | ATGGATTATGGGAAAATGCTC |
| prDH0615 | To screen for T4 *pseT* knockout mutant | AACAGTTCATCCTAATTGGCCT |
| prDH0616 | To screen for T4 *rnlA* mutants | ATCTGCATTAATTCATTCTGCG |
| prDH0617 | To screen for T4 *cef* knockout mutant | CATTCAGCTTAAGCATGGTGTT |
| prDH0618 | To screen for T4 *nrdC.1* knockout mutant | GAATTATGGTCTTTCACTGTTTGAA |
| prDH0619 | To screen for T4 *57B* knockout mutant | TCAAGATCATGAAATGGATGC |
| prDH0620 | To screen for Bas37 *0053* and *0055* knockout mutant | CAGACCGTTACATTGATGCA |
| prDH0621 | To screen for Bas37 *0119* knockout mutant | GGGTAGAAGATGGCGTTAAGTTA |
| prDH0622 | To screen for Bas37 *0130* knockout mutant | CATCGCTATCGTAATATGGAATTAG |
| prDH0623 | To screen for Bas37 *0154* knockout mutant | CTCGACGTGGTATCATGATTTATAC |
| prDH0624 | To screen for Bas37 *0203* knockout mutant | TGGTGACAAGAAAGGCTCATA |
| prDH0625 | To screen for Bas37 *0239 and 0240* knockout mutant | AGGTTGTGATTAACATGTTCACTG |
| prDH0638 | To screen for Bas37 *0033* knockout mutant | TAAAGAATTCGCCGAAGTAGC |
| prDH0650 | To screen for Bas37 *0260* knockout mutant | AGGTGCTACACACGATTTTGAT |
| prDH0626 | To screen for Bas54 *0063* knockout mutant | AAAGTTAGAGGGCAAGCAAG |
| prDH0628 | To screen for Bas54 *0140* knockout mutant | GTCAAGCTGGATTAATCAAGAAG |
| pAH160-P5univ-Fw-PAGE | P7 primer for the second PCR during library preparation of all HIDEN-SEQ libraries | AATGATACGGCGACCACCGAGATCTACACTCTTTCCCTACACGACGCTCTTCCGATCTcgtagaccggggacttatc |
| pAH160-Biot-Fw | Biotinylated primer for the first PCR during library preparation of HIDEN-SEQ libraries with AcrVIA1 as selection marker | ccgggagtctgcgaagttag |
| pDH1-Biot-Fw | Biotinylated primer for the first PCR during library preparation of HIDEN-SEQ libraries with AIcrVIA3 as selection marker | gccgaggagtataatctgaaagttg |
| AcrVIA1 | Synthesized DNA sequence of AcrVIA1 for cloning pAH160_Tn_AcrVIA1_v2 | GCTGTTTTGTGGAATATCTACCGACTGGAAACAGCGATGTTCCGGCCTGCCAGACTGGACGGACATAACAGGTTGGCTGATAAGTCCCCGGTCTGCGGGACTCTGGGGTTCGAAATGACTCGAGATTGACAGCTAGCTCAGTCCTAGGTATAATGCTAGCGGATCCTCTAGATTTTTTAAGAAGGAGATATACATATGATTTACTACATTAAAGATTTGAAGGTAAAGGGGAAAATTTTCGAGAATCTCATGAACAAAGAGGCCGTCGAAGGCCTGATAACCTTTCTGAAGAAGGCAGAGTTCGAGATATATTCTCGGGAGAACTACTCCAAATATAATAAATGGTTTGAGATGTGGAAGTCGCCTACCAGCTCACTGGTCTTCTGGAAGAATTACTCATTTCGTTGTCACCTGTTGTTCGTGATCGAGAAAGACGGTGAATGTTTAGGTATTCCAGCGTCAGTCTTTGAAAGCGTATTGCAGATCTATCTGGCTGATCCCTTTGCTCCGGACACTAAGGAATTGTTCGTGGAAGTATGTAACTTATACGAGTGTCTGGCAGACGTGACGGTAGTTGAGCACTTTGAAGCAGAGGAGTCTGCTTGGCATAAGCTGACGCACAACGAGACAGAAGTCTCAAAGCGCGTCTACTCTAAGGACGATGACGAGTTGCTGAAATATATACCTGAGTTCTTAGACACAATTGCCACGAACAAGAAGTCTCAAAAGTACAATCAAATTCAAGGTAAAATACAGGAGATAAACAAGGAGATCGCCACTTTATACGAGTCTAGTGAGGACTACATATTCACGGAGTATGTATCAAACCTCTACCGGGAGTCTGCGAAGTTAGAACAACACTCGAAACAAATACTGAAGGAAGAGTTAAACTAACTGATTAGAAACAGAGACCTCGTTTACCTATCGGTCTCATGCTGATCTGATAGAGAAGGGTTTGCTCGTAGACCGGGGACTTATCAGCCAACCTGTTATGTGGCGCAGCTAGTTTGTTATCAGAATCGCAGACGACAAGCTGACGACCGCGCAAGTGGCACTTTTCGGG |
| transcriptional terminator | Synthesized DNA sequence of transcriptional terminator for cloning pAH160_Tn_AcrVIA1_v3 | ATACTGAAGGAAGAGTTAAACTAACTGATTAGAAACAGAGACCTCGTTTACCTATCGGTCTCATGCTGTCGGGGAAATGTTCTAGAGGCATCAAATAAAACGAAAGGCTCAGTCGAAAGACTGGGCCTTTCGTTTTATCTGTTGTTTGTCGGTGAACGCTCTCCTGAGTAGGACAAATCCGCCGCCCTAGACCTAGGGTACGGGTTTTGCATCTGATAGAGAAGGGTTTGCTCGTAGACCGGGGACTTATCAGCCAACCTGTTATGTGGCGCAGGTAAACCAGCAATAGACATAAGCGGGAACCGACGACAAGCTGACGACCGCGCAAGTGGCACTTTTCGGG |
| sup-tRNAs | Synthesized DNA sequence of *sup-tRNAs* for cloning pAH160_Tn_sup-tRNAs_LE392_v2 | GCTGTTTTGTGGAATATCTACCGACTGGAAACAGCGATGTTCCGGCCTGCCAGACTGGACGGACATAACAGGTTGGCTGATAAGTCCCCGGTCTGCGGGACTCTGGGGTTCGAAATGACCTAACCAAACAGTCACTTTCGAGCAATTTTCCTTGAAAAAGAGGTTGACGCTGCAAGGCTCTATACGCATAATGCGCCCCGCAACGCCGATAAGGTATCGCGAAAAAAAAGATGGTGGGGTTCCCGAGCGGCCAAAGGGAGCAGACTCTAAATCTGCCGTCATCGACTTCGAAGGTTCGAATCCTTCCCCCACCACCATCGAAGAAACAATCTATTTATTCAAGACGCTTACCTTGTAAGTGCACCCAGTTGGGGTATCGCCAAGCGGTAAGGCACCGGATTCTAATTCCGGCATTCCGAGGTTCGAATCCTCGTACCCCAGCCACATTAAAAAAGCTCGCTTCGGCGAGCTTTTTGCTTTTCTGCGTTCATTCAATGTCGAATGCGATGTTGAAACAGAGACCTCGTTTACCTATCGGTCTCATGCTGATCTGATAGAGAAGGGTTTGCTCGTAGACCGGGGACTTATCAGCCAACCTGTTATGTGGCGCAGGTAAACCAGCAATAGACATAAGCGGGAACCGACGACAAGCTGACGACCGCGCAAGTGGCACTTTTCGGG |
| AIcrVIA3 | Synthesized DNA sequence of AIcrVIA3 for cloning pAH160_Tn_AIcrVIA3_v2 | GCTGTTTTGTGGAATATCTACCGACTGGAAACAGCGATGTTCCGGCCTGCCAGACTGGACGGACATAACAGGTTGGCTGATAAGTCCCCGGTCTGCGGGACTCTGGGGTTCGAAATGACTCGAGATTGACAGCTAGCTCAGTCCTAGGTATAATGCTAGCGGATCCTCTAGATTTTTTAAGAAGGAGATATACATATGGCACCTAAAAAATACGTCGTCACCGTAACAATTCCTGTGACAGATGCAGACCTGACGGTTTTTCTGGTTGTCGATATAATAGTGTATGCGGAGAAACTGGGCGGTACGGTGACAATTACAGCCGTGAAATCCGAAAATGACTCTTATAGCGTTACTCTGGAAGACCTGGATAAAGCGGCCGAAGAATTAGAAAAAGTCGGTGGTAGCATTGTATTGACTGTTACATTTGATAATAAAGAGAAAGCAGAGAAAGTGGCAGAGTTTGCTTACCTGAAAGCCGAGGAGTATAATCTGAAAGTTGATGTTGAAGTTAAAGAGGAACTGTAACTGATTAGAAACAGAGACCTCGTTTACCTATCGGTCTCATGCTGATCTGATAGAGAAGGGTTTGCTCGTAGACCGGGGACTTATCAGCCAACCTGTTATGTGGCGCAGCTAGTTTGTTATCAGAATCGCAGACGACAAGCTGACGACCGCGCAAGTGGCACTTTTCGGG |
| LseCas13a fragment I | Synthesized DNA sequence of LseCas13a for cloning pAH221_LseCas13a | CTAATGCGCTGTTAATCACTTTACTTTTATCTAATCTAGACATCATTAATTCCTAATTTTTGTTGACACTCTATCGTTGATAGAGTTATTTTACCACTCCCTATCAGTGATAGAGAAAAGAATTCAAAAGATCTAAAGAGGAGAAAGGATCTTACATGTGGATTTCCATTAAGACACTGATTCATCACCTGGGAGTTCTGTTTTTTTGTGATTACATGTATAATCGTCGGGAAAAGAAAATTATTGAAGTTAAAACCATGCGCATTACGAAAGTAGAAGTTGATAGAAAAAAAGTTCTTATTTCCCGCGATAAAAACGGTGGCAAGTTAGTGTACGAAAATGAAATGCAGGATAATACTGAACAGATCATGCACCATAAAAAATCATCATTCTACAAATCTGTGGTTAACAAAACAATCTGTCGACCAGAGCAGAAACAGATGAAGAAGCTGGTGCATGGATTGTTACAGGAAAATTCACAGGAAAAAATCAAAGTTAGTGATGTTACAAAGCTCAATATTTCCAATTTCCTGAATCACCGCTTCAAGAAGAGTTTATACTATTTCCCCGAGAATTCGCCTGATAAGTCCGAGGAATATCGCATCGAGATCAATTTGAGCCAACTGTTAGAGGATTCTCTGAAAAAACAGCAAGGTACATTTATTTGTTGGGAATCCTTTAGTAAAGATATGGAACTGTATATTAACTGGGCCGAAAATTATATTAGTTCAAAGACGAAACTGATTAAAAAAAGTATCCGCAACAATCGGATACAGAGCACTGAATCGAGATCCGGGCAGCTCATGGATAGATATATGAAAGACATACTGAATAAGAACAAACCTTTTGATATCCAGTCCGTCTCAGAAAAATATCAATTAGAAAAACTGACCTCTGCTTTGAAGGCAACCTTTAAAGAAGCTAAAAAAAACGACAAAGAAATTAATTACAAACTCAAATCAACGCTGCAGAACCATGAACGGCAGATTATAGAGGAACTTAAAGAGAACTCGGAGCTTAACCAGTTTAACATCGAAATACGCAAACATCTGGAAACCTACTTCCCAATTAAAAAAACCAATCGTAAAGTAGGAGATATTCGAAATCTTGAAATCGGCGAAATCCAGAAAATCGTCAATCATCGTCTTAAGAACAAAATTGTGCAGCGCATCCTGCAGGAGGGCAAATTGGCGTCTTATGAAATTGAAAGTACCGTGAATTCCAATAG |
| LseCas13a fragment II | Synthesized DNA sequence of LseCas13a for cloning pAH221_LseCas13a | AAATTGAAAGTACCGTGAATTCCAATAGCCTCCAGAAAATTAAAATTGAAGAGGCGTTCGCCCTTAAATTTATTAATGCATGCTTATTCGCGTCCAACAATCTGCGCAACATGGTATATCCTGTTTGTAAAAAAGACATTCTTATGATTGGTGAATTCAAGAATTCTTTTAAGGAAATTAAACATAAAAAATTTATTCGACAGTGGTCCCAGTTCTTTTCCCAAGAAATTACCGTGGACGACATTGAGCTTGCTAGCTGGGGTCTGCGTGGTGCTATCGCACCAATTAGAAATGAGATCATTCACCTCAAGAAACACTCATGGAAAAAATTCTTCAACAACCCAACATTCAAGGTGAAAAAGTCCAAAATCATAAATGGGAAAACCAAAGATGTGACCAGCGAGTTTCTCTATAAAGAGACCCTGTTCAAGGATTATTTCTATAGCGAGCTGGACAGCGTCCCAGAGCTCATTATTAACAAGATGGAGTCATCAAAAATATTAGATTATTACAGCTCTGATCAGCTCAATCAGGTGTTTACCATTCCAAATTTCGAACTGTCACTTCTTACCTCAGCTGTTCCGTTTGCACCATCGTTTAAACGTGTTTATCTGAAAGGATTTGATTATCAGAATCAGGATGAGGCACAACCGGACTACAATTTGAAATTAAATATATACAATGAAAAAGCCTTTAATAGTGAAGCCTTTCAAGCACAATATTCACTCTTTAAAATGGTGTATTATCAGGTGTTTCTGCCGCAATTTACAACGAATAATGATCTGTTTAAATCGAGCGTGGATTTTATCCTGACACTGAATAAGGAACGCAAGGGATATGCCAAAGCGTTTCAAGATATTAGAAAAATGAACAAAGATGAAAAACCTTCAGAGTACATGTCATATATTCAGAGTCAACTGATGCTGTATCAAAAAAAGCAGGAAGAAAAGGAGAAAATTAATCATTTTGAGAAATTTATTAACCAGGTGTTTATAAAAGGTTTTAATTCCTTTATTGAAAAGAATCGTCTGACCTATATATGTCATCCTACCAAAAATACGGTACCAGAAAACGATAATATCGAAATTCCGTTTCATACGGACATGGATGACTCAAACATTGCTTTTTGGTTAATGTGCAAACTGCTTGATGCGAAACAGCTGAGTGAGCTCCGTAACGAGATGATCAAATTCTCATGTTCCTTGCAGTCCACCGAGGAAATTAGCACGTTCACTAAGGCCCGCGAGGTCATTGGATTGGCATTACTCAACGGTGAAAAAGGCTGCAATGATTGGAAGGAATTATTTGATGATAAAGAAGCCTGGAAAAAAAATATGTCTCTGTATGTATCTGAAGAGCTGTTACAGTC |
| LseCas13a fragment III | Synthesized DNA sequence of LseCas13a for cloning pAH221_LseCas13a | GTATGTATCTGAAGAGCTGTTACAGTCCTTGCCCTACACGCAGGAAGATGGTCAGACGCCCGTTATTAATCGCTCTATCGACCTCGTTAAGAAATACGGCACAGAAACCATTTTAGAAAAATTGTTCTCATCTTCGGACGACTACAAAGTCTCTGCGAAAGATATTGCAAAGTTGCATGAGTATGACGTAACAGAAAAAATTGCACAGCAGGAAAGCTTGCATAAACAGTGGATCGAAAAACCTGGACTGGCACGTGATTCAGCGTGGACAAAGAAATATCAAAATGTCATAAATGATATTTCTAATTATCAATGGGCCAAAACTAAAGTAGAACTTACCCAAGTTCGCCATCTTCATCAGTTAACGATTGATTTGTTGAGTCGTCTGGCGGGCTACATGTCTATTGCAGATCGTGACTTTCAGTTTAGCTCCAATTATATCTTGGAAAGAGAGAATAGCGAGTACCGTGTTACCTCCTGGATTCTGCTGTCTGAGAACAAGAATAAGAACAAGTACAACGACTACGAGCTGTATAATCTTAAAAACGCGTCCATCAAAGTCAGTAGCAAGAATGATCCGCAACTTAAAGTTGATCTGAAGCAGCTGCGACTGACACTCGAATATTTAGAGCTGTTTGATAACAGATTAAAAGAAAAGCGTAATAATATTAGCCACTTCAATTACTTAAATGGACAGTTAGGTAACAGTATATTGGAATTATTCGACGACGCCCGCGATGTACTGAGCTATGATCGTAAGCTGAAGAATGCCGTTTCCAAATCGCTGAAAGAAATTTTATCCTCTCATGGCATGGAAGTTACATTTAAGCCCTTATACCAAACCAACCATCATCTCAAAATCGACAAATTACAACCTAAGAAAATCCATCACCTGGGTGAGAAATCAACTGTATCTTCGAATCAGGTATCTAACGAGTACTGTCAACTTGTCAGAACCCTGTTAACAATGAAGTAATGAATCCCTCGGTACCAAAGACGAACAATAAGACGCTGAAAAGCGTCTTTTTTCGTTTTGGTCCGCTGAGCAGTTACAGAGATGTTACGAACCACTAGTGCACTGCAGTACACCTTTGGTCGAAAAAAAAAGCCCGCACTGTCAGGTGCGGGCTTTTTTCTGTGTTTCCACCAATAAAAAACGCCCGGCGGCAACCGAGCGTTCTGAACAAATCCAGATGGAGTTCTGAGGTCATTACTGGATCTATCACGCTAGTTTGTTATCAGAATCGCAG |
| crRNA LseCas13a | Synthesized DNA sequence of LseCas13a crRNA cassette for cloning the entry vector pAH218_LseCas13a | GGAATTCTTGACAGCTAGCTCAGTCCTAGGTATAATACTAGTGTAAGAGACTACCTCTATATGAAAGAGGACTAAAACAGAGACCTCGTTTACCTATCGGTCTCATGCTTGGGCCCGAACAAAAACTCATCTCAGAAGAGGATCTGAATAGCGCCGTCGACCATCATCATCATCATCATTGAGTTTAAACGCTCTCCAGCTTGGGTGTTTTGGCGGATGAGAGAAGATTTTCAGCCTGATACAGATTAAATCAGAACGCAGAAGCGGTCTGATAAAACAGAATTTGCCTGGCGGCAGTAGCGCGGTGGTCCCACCTGACCCCATGCCGAACTCAGAAGTGAAACGCCGTAGCGCCGATGGTAGTGTGGGGTCTGCCCATGCGAGAGTAGGCAACTGCCAGGCATCAAATAAAACGAAAGG |
